# Bayesian parameter inference for epithelial mechanics

**DOI:** 10.1101/2024.04.24.590933

**Authors:** Xin Yan, Goshi Ogita, Shuji Ishihara, Kaoru Sugimura

**Author notes:** Correspondence (GO), (KS). Joint first authors (Xin Yan, Goshi Ogita).

## Abstract

Cell-based mechanical models, such as the Cell Vertex Model (CVM), have proven useful for studying the mechanical control of epithelial tissue dynamics. We recently developed a statistical method called image-based parameter inference for formulating CVM model functions and estimating their parameters from image data of epithelial tissues. In this study, we employed Bayesian statistics to improve the utility and flexibility of image-based parameter inference. Tests on synthetic data confirmed that both our non-hierarchical and hierarchical Bayesian models provide accurate estimates of model parameters. By applying this method to *Drosophila* wings, we demonstrated that the reliability of parameter estimation is closely linked to the mechanical anisotropies present in the tissue. Moreover, we revealed that the cortical elasticity term is dispensable for explaining force-shape correlations *in vivo*. We anticipate that the flexibility of the Bayesian statistical framework will facilitate the integration of various types of information, thereby contributing to the quantitative dissection of the mechanical control of tissue dynamics.

## 1 Introduction

Mathematical modeling provides means for testing hypotheses that are experimentally inaccessible, deciphering underlying mechanisms, and guiding new research directions through testable predictions [1–3]. These advantages make it indispensable for understanding complex biological phenomena, such as tissue development and repair. Mathematical modeling typically involves three key steps: model construction, parameter estimation, and model selection [4, 5]. First, model equations are formulated to represent the biological process of interest, often resulting in multiple candidate models. Subsequently, parameter estimation is carried out to determine the parameter values that best recapitulate experimental data. Assessing the uncertainty associated with parameter estimation is important, especially when making decisions based on theoretical predictions [6–8]. Finally, the most appropriate model is selected from the candidate models based on statistical criteria that balance the goodness-of-fit and model complexity. Despite its importance, a statistical framework to evaluate parameter estimation uncertainty and model goodness is yet to be fully established in developmental biology [8, 9].

Various modeling frameworks have been employed to investigate the physical mechanisms by which mechanics control tissue dynamics [10–13]. Among them, the cell vertex model (CVM) is one of the most frequently used models for studying epithelial mechanics [14–16]. The CVM treats epithelial tissue as a tile of polygons, and computes the displacement and reconnection of cell vertices via the minimization of virtual work, by which the balance between junction tensions and cell pressure is considered (Fig. 1A-C). In the CVM, cellular forces, such as junction tension and cell pressure, are dependent on the morphological characteristics of cells (*e*.*g*., junction length and cell area), for which specific forms of model functions and parameter values are assumed. These model functions and parameters must be determined based on experimental data to ensure the validity of the modeling; however, most existing approaches do not fully leverage the quantitative information embedded in experimental data.

**Figure 1:**
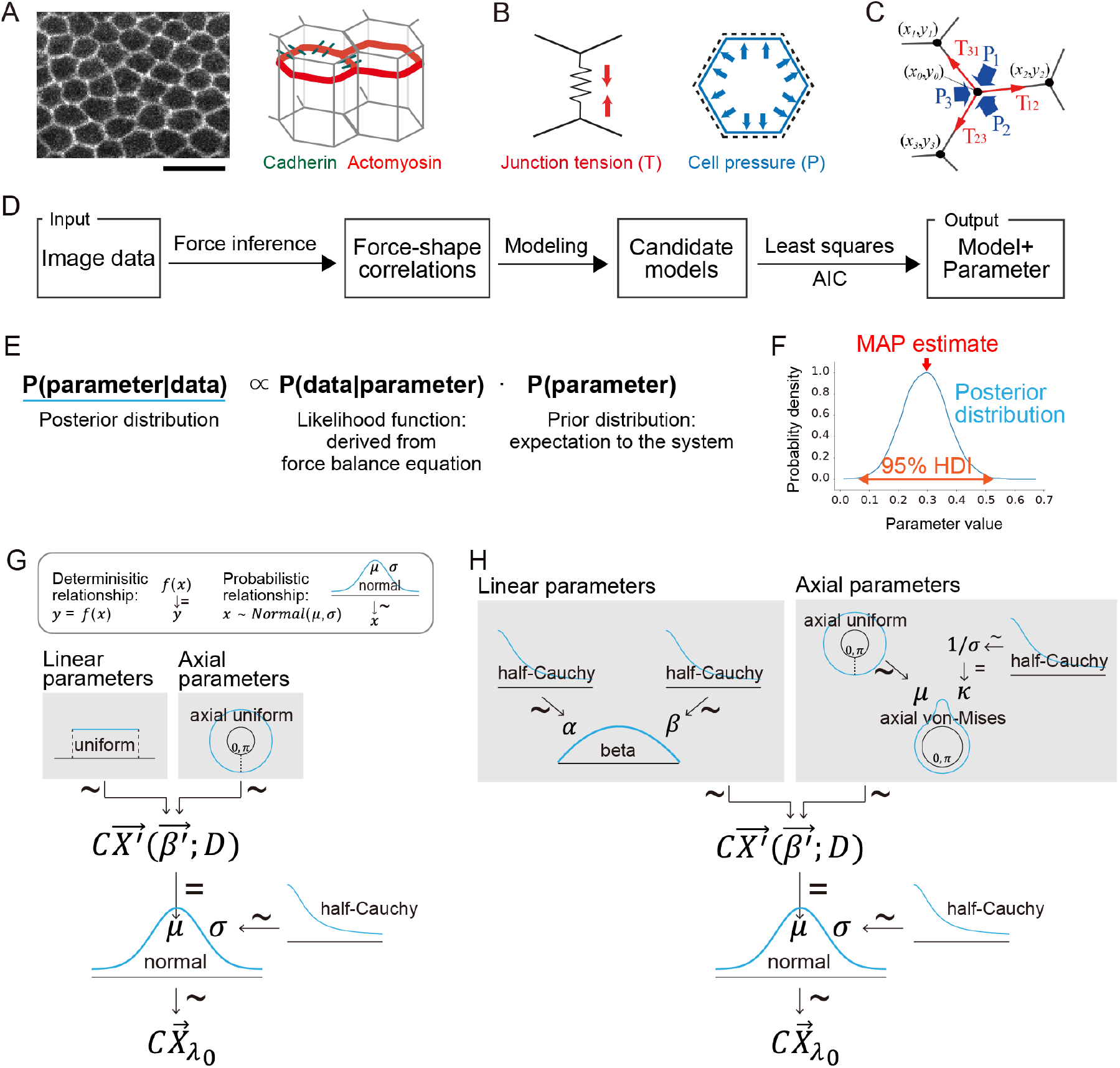
Bayesian parameter inference for epithelial mechanics. (A) The structure of the monolayer epithelium. Left: The top view of the *Drosophila* pupal wing epithelium, in which the adherens junction (AJ) is labeled with DE-cadherin-GFP. Right: Epithelial cells are connected to each other via cell adhesion molecules such as cadherin (green). Actomyosin cables (red) run along the cell cortex in the AJ plane. Scale bar: 10 *µm*. Forces acting along the AJ plane. Left: Junction tension. Right: Cell pressure. Balance of junction tensions (T) and cell pressures (P) acting on the cell vertex. Black dots indicate cell vertices. (D) Flowchart of the method for estimating mechanical parameters from image data, in which the cell contour is visualized [27]. (E) According to Bayes’ theorem, the posterior distribution *P*(parameter|data) is calculated from the likelihood *P*(data|parameter) and the prior distribution *P*(parameter). It provides an estimate of a parameter and uncertainty associated with estimation. In the image-based parameter inference, the likelihood function is derived from the force balance equation at cell vertices. (F) An example of the posterior distribution obtained by MCMC sampling. The red arrow indicates the maximum a posteriori (MAP) estimate, while the orange double-headed arrow represents the 95% highest density interval (HDI), which quantifies the estimation uncertainty. (G, H) The Kruschke style diagram [29] of the non-hierarchical and hierarchical Bayesian models used in this study (G: non-hierarchical, H: hierarchical). The characters represent the parameters or variables whose dependencies are indicated by arrows. The symbols *∼* and = next to the arrows respectively indicate probabilistic and deterministic dependency. See section 2.4 for details about the formulation of Bayesian models. (A)–(D) are adapted from [27].

With some recent exceptions [17–21], many previous studies using the CVM have relied on standard model functions that qualitatively represent epithelial mechanics [15, 16]. Parameter values are determined by iteratively running numerical simulations with different parameter sets and fitting the simulation results to experimental observations [7, 22–26]. Summary statistics, such as the polygonal distribution of cells, are used for this fitting process, making the comparison between simulation and experimental data indirect. Moreover, information criteria are often underutilized for comparing multiple candidate models.

To address these technical challenges concerning the CVM, we recently developed a data-driven, statistical method to deduce model functions and estimate their parameters directly from experimental data [27] (Fig. 1D; explained in detail in Methods). First, we formulate model functions (tension and pressure functions) based on correlations between force and cell shape obtained from image data of epithelial tissue. Substitution of the model functions into force-balance equations at the cell vertex leads to an equation with respect to the parameters of the model. Solving this equation using a least-squares method allows for the estimation of parameter values, without performing numerical simulations. Finally, Akaike Information Criterion (AIC) [28] is employed for selecting the most suitable model among candidates. Tests using synthetic and *in vivo* data have validated the accuracy and robustness of parameter estimation and model selection [27]. Overall, our method offers a simple, efficient, and data-driven approach for cell-based modeling in epithelial mechanics.

Nevertheless, the current method has some limitations. It provides only point estimates of parameters, leaving the uncertainty in estimation unaddressed. In addition, the point estimation makes it difficult to identify representative models and parameters for groups categorized by tissue type, developmental stage, genotype, and so on. Furthermore, certain model functions result in physically unrealistic estimates, likely due to multicollinearity. Bayesian statistics have been known to offer solutions for overcoming related limitations [29, 30]. The uncertainty in parameter estimation can be directly quantified from the posterior distribution. Hierarchical Bayesian modeling enables the integration of multi-layered information, such as groups and their individual samples. Using informative priors to restrict the parameter space may extend the range of model functions that can be analyzed.

In this study, we aimed to improve the image-based parameter inference using Bayesian statistics. We developed both non-hierarchical and hierarchical Bayesian models and evaluated their accuracy in parameter estimation. With these methods, we explored how uncertainty in parameter estimation correlates with developmental changes in tissue mechanics, and examined whether mathematical representation of the cortical elasticity is appropriate for modeling epithelial tissues. We foresee that the flexibility in Bayesian statistical framework will help integrating various types of information such as time and space, thereby facilitating the quantitative dissection of mechanical control of tissue dynamics.

## 2 Methods

In this study, we reformulated image-based parameter inference [27] using Bayesian statistics. The original method and the new Bayesian method follow the same procedures until constructing candidate models (section 2.1) and writing force-balance equations at cell vertices (section 2.2). After that, the former uses the least-squares method for parameter estimation (section 2.3), while the latter formulates the problem using Bayes’ theorem (section 2.4).

### 2.1 Mechanical model for epithelial tissue

In the monolayer epithelium, actomyosin cables running along the plane of adherens junction (AJ) generate junction tension, while cell pressure counteracts junction tension to maintain the cell area (Fig. 1A–C). In the CVM, the partial differentiation of virtual work 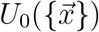 by the length of the junction between the *i* -th and *j* -the cell, *l*_*ij*_, leads to the junction tension *T*_*ij*_ as *T*_*ij*_ *≡* ∂*U*_0_*/*∂*l*_*ij*_, while the partial differentiation by the area of the *i*-th cell, *A*_*i*_, leads to the cell pressure *P*_*i*_ as *P*_*i*_ *≡ −* ∂*U*_0_*/*∂*A*_*i*_. The junction tension and the cell pressure are determined by model equations that include parameters representing cell mechanical properties, such as the elastic coefficient of cell.

We have established data-driven formulation of tension and pressure equations in the CVM (Fig. 1D) [27]. Briefly, we first employed Bayesian force inference [31] to quantify the relative values of junction tension and pressure difference among cells from image data of epithelial tissues. Subsequently, we plotted the inferred forces against the morphological characteristics of cells, such as the junction length and the cell area. Through this analysis of force-shape correlation, we formulated candidate model functions for tension and pressure (for detailed information, see [27]).

#### 2.1.1 Model equations for junction tension

The function for junction tension is of the following form:

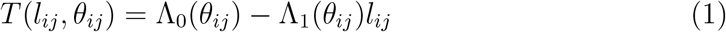

where *θ*_*ij*_ and *l*_*ij*_ are the direction and length of junction between *i* -th and *j* -th cells. The anisotropic line tension Λ_0_(*θ*) (Fig. 2A) and the anisotropic spring constant Λ_1_(*θ*) (Fig. 2A) are defined as bellow:

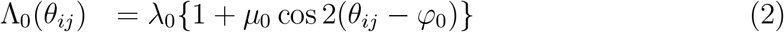

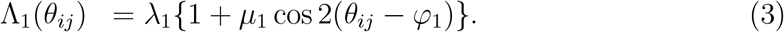

*λ*_0_ and *λ*_1_ represent the magnitude of Λ_0_(*θ*_*ij*_) and Λ_1_(*θ*_*ij*_), respectively. *µ*_0_ and *µ*_1_ denote the magnitudes of their orientation-dependent (anisotropic) parts, which peak at *φ*_0_ and *φ*_1_. *λ*_0_, *λ*_1_, *µ*_0_ and *µ*_1_ are the linear parameters and *φ*_0_ and *φ*_1_ are the axial parameters with interval [0, *π*].

**Figure 2:**
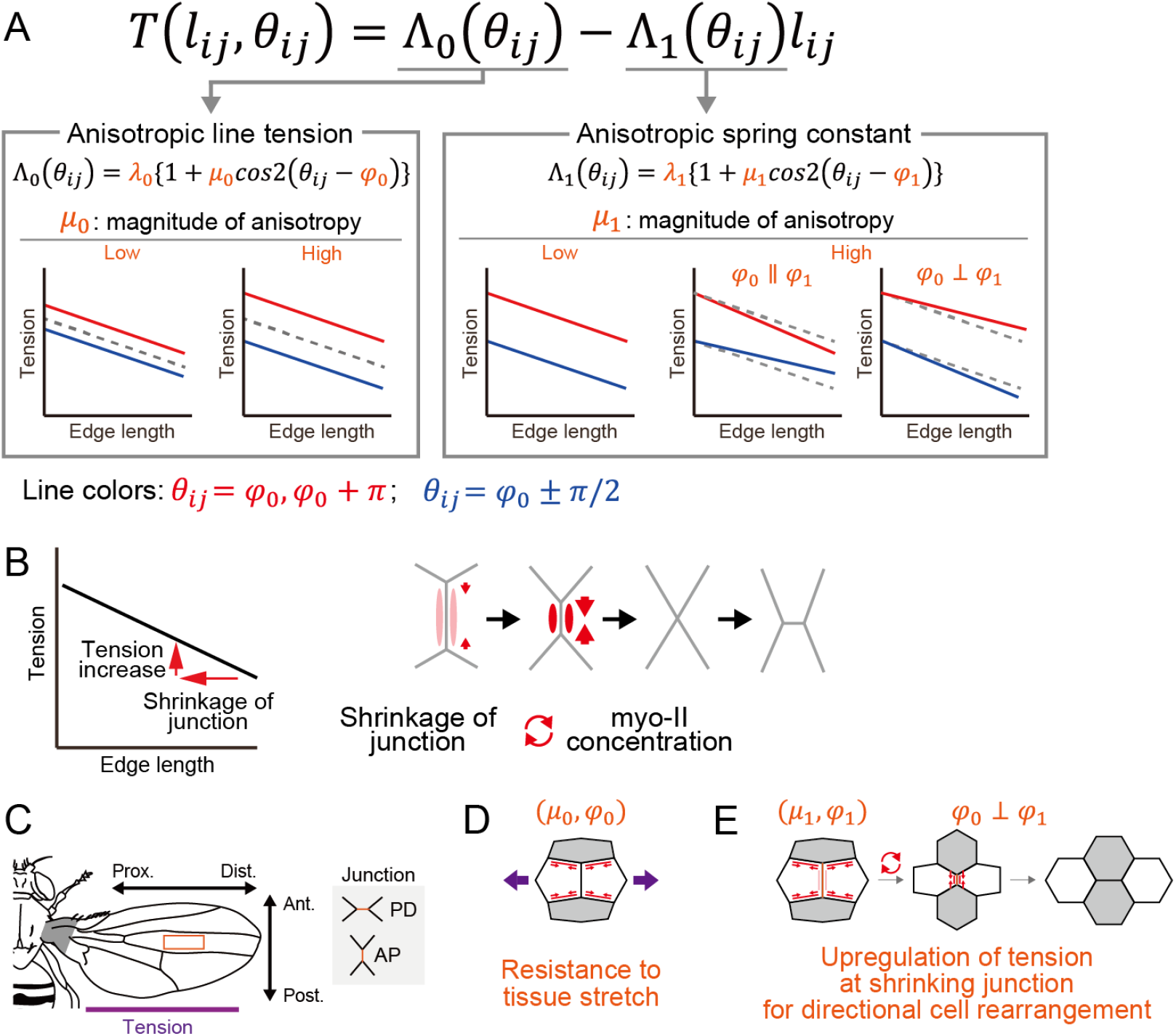
The anisotropic spring model for junction tension. (A) The anisotropic spring model consists of two terms, Λ_0_(*θ*_*ij*_) and Λ_1_(*θ*_*ij*_)*l*_*ij*_ (section 2.1.1) [27]. Λ_0_(*θ*_*ij*_) represents line tension, with the magnitude and orientation of the anisotropy determined by *µ*_0_ and *φ*_0_, respectively. Λ_1_(*θ*_*ij*_) represents junction length-dependent junction tension, with its magnitude and orientation of anisotropy determined by *µ*_1_ and *φ*_1_, respectively. Schematics show how the values of *µ*_0_ and *µ*_1_ alter the dependence of tension on junction length and orientation, in which line colors denote the orientation of junctions. The right-most plot corresponds to the pupal wing, where the red and blue lines represent junctions along the proximal-distal (PD) and anterior-posterior (AP) axes, respectively. (B) Schematics illustrating the feedback between junction shrinkage and myosin-II (myo-II)-generated junction tension. When the junction tension is inversely related to the junction length, shrinkage of a junction (indicated by the left-pointing arrow) leads to an increase in junction tension (indicated by the upward arrow), which in turn shortens the junction. This feedback is associated with the accumulation of myo-II at shrinking junctions. Red arrows represent junction tension, while the color intensity indicates myo-II concentration along the junctions. (C) Schematic of the adult fly. Orange rectangle indicates the wing region studied in this paper. Purple line indicates the orientation of tissue tension. Schematics of junction are shown in the right. The hinge is shaded gray. (D, E) Schematics illustrating the biological functions of Λ_0_(*θ*_*ij*_) and Λ_1_(*θ*_*ij*_) during the development of the *Drosophila* pupal wing [27]. Red arrows represent the junction tension, and purple double-sided arrows indicate the extrinsic pulling force from the hinge. (D) Λ_0_(*θ*_*ij*_) contributes to resistance against the extrinsic pulling force from the hinge. (E) Λ_1_(*θ*_*ij*_) facilitates the shrinkage of AP junctions by upregulating the junction tension through the feedback depicted in (B). The myo-II enrichment along the remodeling junctions is indicated by the change in line colors from pale pink to red along the vertical shrinking junction. (B) and (C)–(E) are adapted from [66] and [27], respectively.

The anisotropic spring constant, Λ_1_(*θ*_*ij*_), represents feedback between myosin-II (myo-II)-generated junction tension and junction shrinkage (Fig. 2B). The sign of Λ_1_(*θ*_*ij*_) indicates whether the feedback is positive (*i*.*e*., accelerating junction shortening/lengthening) or negative (*i*.*e*., stabilizing the junction length). This feedback between tension/myo-II and junction shrinkage is supported by experimental observations that myo-II is getting concentrated on shrinking junctions during cell rearrangement [17, 32, 33]. Related but more complex models representing the negative dependency of junction tension on junction length or strain have also been proposed [17, 18, 21].

Our previous study identified distinctive biological functions of Λ_0_(*θ*_*ij*_) and Λ_1_(*θ*_*ij*_) during the development of the *Drosophila* pupal wing (Fig. 2C–E) [27]. Specifically, Λ_0_(*θ*_*ij*_) contributes to resistance against the extrinsic pulling force from the hinge. This is supported by observations that *µ*_0_ increases following the onset of hinge constriction at 15 hours after puparium formation (h APF), which exerts a pulling force along the proximal-distal (PD) axis, and that *φ*_0_ aligns with the PD axis (Fig. 2C, D) [34–38]. On the other hand, *φ*_1_ is aligned with the anterior-posterior (AP) axis and Λ_1_(*θ*_*ij*_) promotes the shrinkage of AP junctions, running perpendicular to the global tension anisotropy (Fig. 2E) [35, 38, 39].

In section 3.4, we consider the following alternative model incorporating cortical elasticity:

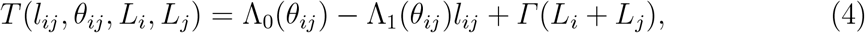

where *Γ* and *L*_*i*_ represent the elastic coefficient of cell cortex and the perimeter of *i* -th cell, respectively.

In addition to the models described above, simpler models were considered as summarized in Fig. S1. Model A is hereafter referred to as the anisotropic spring model. Model E’ corresponds to the standard equation of junction tension commonly used in the CVM [15, 16, 22]:

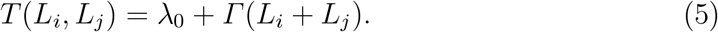

#### 2.1.2 Model equations for cell pressure

The pressure function used in this study is as follows:

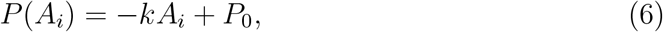

where *A*_*i*_ is the area of *i*-th cell. *k* and *P*_0_ are the elastic coefficient of a cell and the hydro-static pressure, respectively [15, 16, 22]. We have tested alternative pressure functions and demonstrated that the choice of pressure function had a negligible impact on parameter estimation [27].

### 2.2 Force balance equation for epithelial tissue

The parameters involved in tension/pressure functions are estimated by considering the balance of forces acting on the cell vertices [27]. Suppose that the pressure of the *i*-th cell is *P*_*i*_ and the tension of the cell junction between the *i*-th and *j*-th cell is *T*_*ij*_ (Fig. 1B). Under the approximation that epithelial sheet is regarded as a tessellation of polygons, the following equations describe the net forces acting at the 0-th vertex, 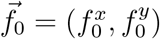 in Fig. 1C [31, 40–42].

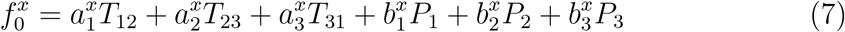

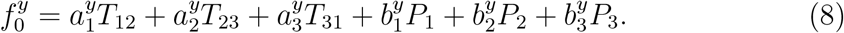

The coefficients are calculated only from the vertex coordinates, such as 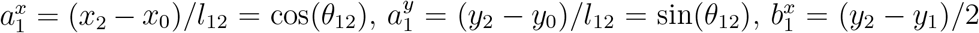 and 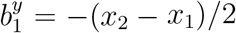, and are measurable from image data.

By repeating this procedure for all vertices in tissue, the following equation is obtained:

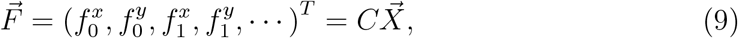

where *C* is an *n × m* matrix composed of coefficients 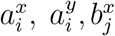 and 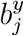 and 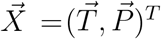 is an *m*-dimensional vector composed of *T*_*ij*_ and *P*_*i*_ [31]. *n* is the number of force balance equations and equals twice the number of vertices. *m* is the sum of the number of junctions and the number of cells, which corresponds to the dimension of the vector 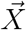. Given that epithelial morphogenesis is assumed to be a quasi-static process, wherein forces acting on the vertices are almost balanced, the following force-balance equation holds:

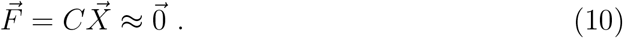

Substitution of the model equations (Eqs. 1 and 6) into Eq. 10 gives an equation of the parameter vector 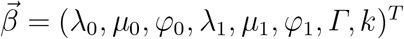

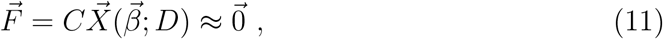

where *D* is the list of cell shape characteristics, namely the junction length *l*_*ij*_, the junction angle *θ*_*ij*_, the cell perimeter length *L*_*i*_, and the cell area *A*_*i*_ quantified from image data. Since the solution to this equation has arbitrariness in the scaling, we chose *λ*_0_ as a scaling factor and fixed it to 1.0. This enables us to decompose 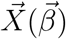 into the constant term derived from *λ*_0_ and the data-dependent term involving the other parameters:

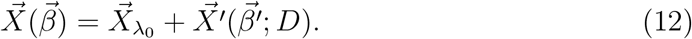

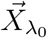 is the *m*-dimensional vector whose component 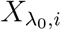 is 1 (*i* = 1, …, *m*_*j*_) or 0 (*i* = *m*_*j*_ + 1, …, *m*), where *m*_*j*_ is the number of junctions.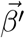 is the vector composed of parameters other than *λ*_0_. 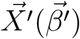 represents the cellular force with the constant term subtracted. Substitution of Eq. 12 into Eq. 11 leads to

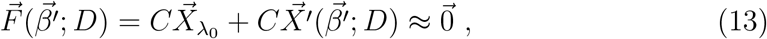

which is the equation of the parameter to be solved.

### 2.3 Parameter estimation by the least-squares method

In this section, we outline the procedure used in [27] to estimate parameters by transforming Eq. 13 into the multiple-linear regression problem and solving it by the least-squares method.

The estimated values of the mechanical parameters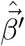 are obtained by minimizing the *L*^2^ norm of 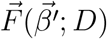 in Eq. 13:

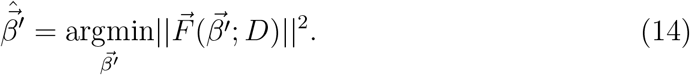

To transform this minimization problem in Eq. 14 into the multivariate regression problem, we rewrite the anisotropic line tension Λ_0_(*θ*_*ij*_) and the anisotropic spring constant of junction Λ_1_(*θ*_*ij*_) as:

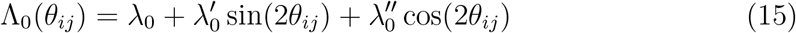

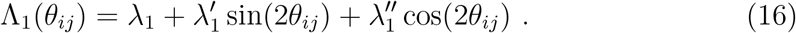

The transformed parameters 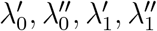 are defined as 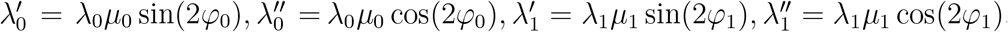. Next, we define the matrices *S*_*T*_ and *S*_*P*_ as follows:

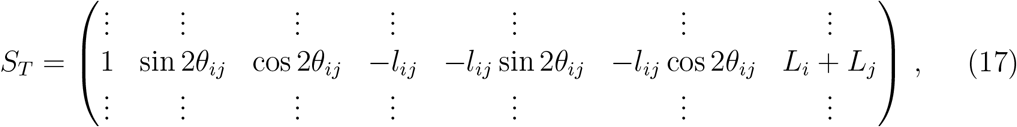

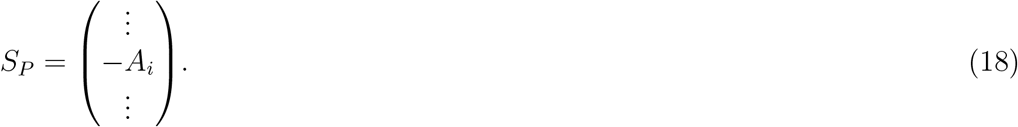

Note that *S*_*T*_ and *S*_*P*_ are calculated based on *D*, the list of cell shape characteristics quantified from image data. Consequently, the vectors 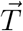 and 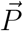 representing junction tension and cellular pressure are given by

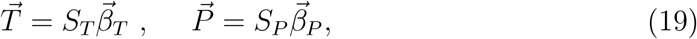

respectively, where 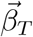 and 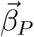 are composed of the transformed mechanical parameters.

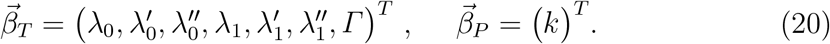

By using Eq. 19, the cellular force vector *X* is denoted as:

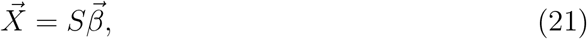

with the matrix *S* and the parameter vector 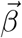 defined as

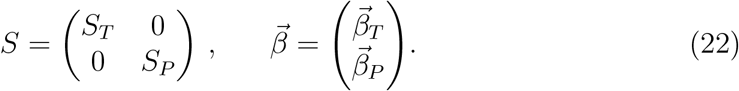

Substituting Eq. 21 into Eq. 13 yields:

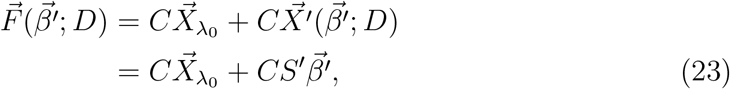

where *S*^*′*^ is obtained by eliminating the first column from *S*. The minimization problem of *L*^2^ norm of 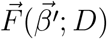 given in Eq. 14 is equivalent to the linear multiple regression problem with 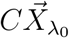 as the observation vector, *−CS*^*′*^ as the design matrix, and 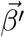 as the parameter vector. This means that the estimation of the parametervector 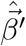 can be obtained analytically as follows:

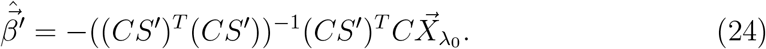

We solved the lest-squares problem using the function **OLS.fit()** in the Python package statsmodels.

When analyzing individual samples separately, we derived *C* and *S* from image data of each sample and then performed parameter estimation. For pooled estimation, where individual samples are aggregated into a single dataset, we combined *C* (or *S*) for image data of all samples in the group before proceeding with parameter estimation.

### 2.4 Parameter estimation by Bayesian statistics

In this study, we constructed both a non-hierarchical Bayesian model and a hierarchical Bayesian model (Fig. 1E–H). The former is designed for estimating parameters of individual samples, while the latter is designed for estimating parameters at both the group level and individual level (Fig. 1G, H) [29].

#### 2.4.1 Non-hierarchical Bayesian model

The posterior distribution of the non-hierarchical Bayesian model, 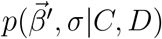, is defined as follows:

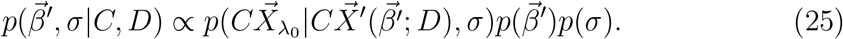

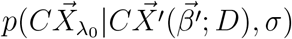 is the likelihood function with a scale parameter *σ* and represents the probability of observing the data at given parameter values 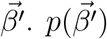 and *p*(*σ*) are the prior distributions for 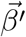 *′* and *σ*, respectively.

The likelihood function is derived from the force-balance equation at cell vertices (section 2.2). We assume that the net forces acting on cell vertices are normally distributed with zero mean and *σ*^2^*I* covariance, where *I* is the *n × n* identity matrix [31]. This assumption is equivalent to 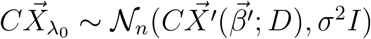 based on Eq. 13.

The likelihood function thus reads:

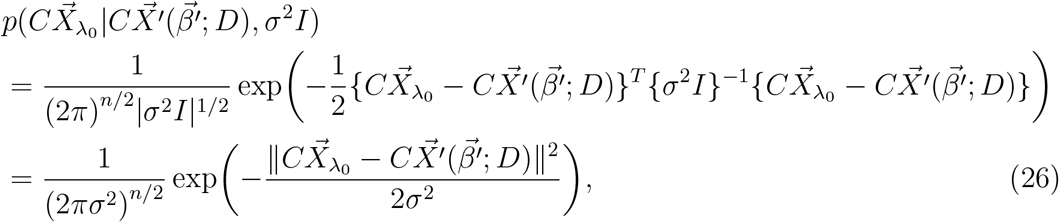

where |*σ*^2^*I*| represents the determinant of *σ*^2^*I*. We employed the following non-informative priors. For the prior distribution of *λ*_1_, *µ*_0_, *µ*_1_ and *k*, we applied a uniform distribution on the interval [0, 1]. For the prior distribution of the elastic coefficient of the cell cortex, *Γ*, we applied a half-Cauchy distribution with a scale of 5. For the prior distribution of *φ*_0_ and *φ*_1_, we applied an axial uniform distribution, which is a uniform distribution on the semi-circle [0, *π*) [43, 44]. We set a half-Cauchy prior with the scale 5 on *σ*. We tested other prior distributions, such as half normal distribution, instead of half Cauchy distribution, and confirmed that the posterior distributions were not significantly affected by the choice of the prior distributions: the maximum difference between estimated parameter values was in relative value of 3.96 ×10^*−*5^ for linear parameters, and in absolute value of 1.26 × 10^*−*5^ radians for axial parameters, in three test data of *Drosophila* epithelial tissues.

#### 2.4.2 Hierarchical Bayesian model

The posterior distribution of a hierarchical Bayesian model, 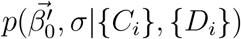, is defined as follows:

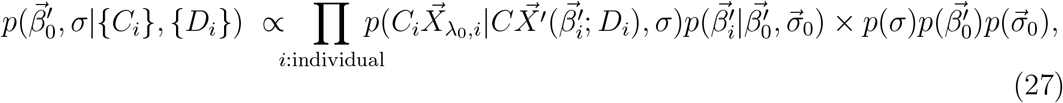

where subscript *i* indicates the index of samples in a group, and {*C*_*i*_} and {*D*_*i*_} are the list of coefficient matrices and cell shape quantities in a group data. The likelihood function 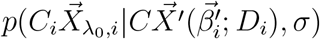 is the same as that in the non-hierarchical Bayesian model. 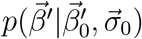 is the group-level distribution of mechanical parameters, which the mechanical parameters of each sample follows. 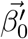 is representative mechanical parameters of a group. 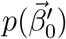 is the prior distribution of the group-level meta parameters 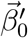.

We employed unimodal distributions for the group-level distribution of the mechanical parameters. For the linear parameters (*λ*_1_, *µ*_0_, *µ*_1_, *k*), we applied the beta distributions. 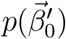 For instance, the group-level distribution of *λ*_1_ is given by:

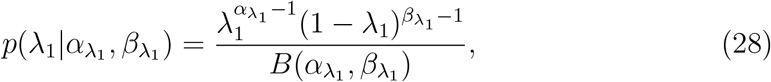

where 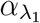 and 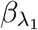 are parameters of the beta distribution and 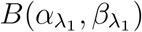 is the beta function. The group-level distributions of the other linear parameters were set in the same way. For the axial parameters (*φ*_0_, *φ*_1_), we utilized the axial von-Mises distribution [44]. For instance, the group-level distribution of *φ*_0_ reads

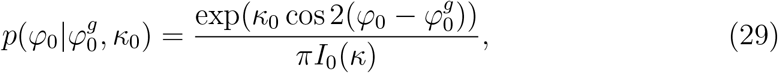

where *I*_0_(*κ*) is the modified Bessel function of order zero. 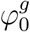 and *κ*_0_ represent the location and concentration parameters of a group, respectively. We applied half-Cauchy priors with the scale 5 on 1*/κ*_0_. The group-level distribution of *φ*_1_ was set following the same procedure.

### 2.5 MCMC Sampling and MAP estimation

We conducted Markov chain Monte Carlo (MCMC) sampling using PyMC3, a python package for Bayesian inference (version 3.11.4; [45]). We applied the ADVI and NUTS algorithms to initialize and sample the posterior distribution [46, 47]. In the NUTS sampler, we set *target accept* = 0.99. For the non-hierarchical Bayesian method, we ran two MCMC chains, each with 15,000 steps, of which the first 5,000 steps were discarded as burn-in. We assessed the quality of sampling using the Gelman-Rubin statistic 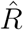 [29, 48, 49] and divergences. If 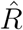 exceeded 1.05 and/or divergences occurred frequently, the entire run was discarded and repeated. The MAP estimate was calculated using the find MAP function in PyMC3, in which the numerical optimization is performed using the CG algorithm. For the hierarchical Bayesian method, we ran six MCMC chains. If most chains converged but one or two chains clearly deviated, we discarded the deviating chains and used the remaining chains for the sampling results. We calculated the MAP estimate from the mode of the posterior distribution sampled by MCMC. The approximate time required for MCMC sampling was 1–2 hours per image data, regardless of using non-hierarchical or hierarchical Bayesian models.

### 2.6 Model selection

For model selection in the Bayesian methods, we employed Widely applicable information criterion (WAIC) and leave-one-out cross-validation (LOOCV) [49–51]. WAIC and LOOCV are respectively calculated by the functions **waic** and **loo** in a python package Arviz [52]. We found that WAIC and LOOCV selected the same model both in the non-hierarchical and hierarchical Bayesian methods in our data sets of *Drosophila* epithelial tissues.

AIC was used for model selection in the original least-squares method [27]. AIC was calculated using **OLS.fit()** in the Python package statsmodels.

### 2.7 Uncertainty quantification

We quantified the uncertainty associated with parameter estimation using the 95% highest posterior density (HDI) interval, within which a parameter value falls with 95% probability in the posterior distribution (Fig. 1F) [29]. The average of the 95% HDIs for each selected model at every developmental stage was calculated as follows: for linear parameters, we normalized the 95% HDIs by the MAP value for each image data. For axial parameters, we first calculated the double lengths of the 95% HDIs using circular statistics [43, 44] for each image data, and then divided these values by 2. We subsequently computed the average of these values for both linear and axial parameters and listed them in Fig. 5A.

### 2.8 Generation of synthetic data set

In this study, we used synthetic data to assess the accuracy of parameter estimation. The synthetic data on the cell configuration was generated through numerical simulation of the CVM as previously described [27]. The positions and connectivity of vertices 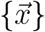 were changed according to the minimization of virtual work 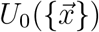 The virtual work of the anisotropic spring model with cortical elasticity (model A’ in Fig. S1) is given as

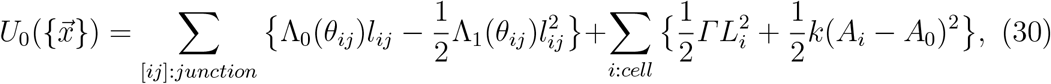

where *A*_0_ is the target area of a cell and set at *A*_0_ = 1.0. We solved the following equation in simulations.

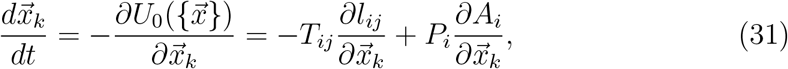

where derivatives of anisotropy coefficients such as 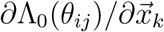 are ignored [53]. The numerical simulations were implemented by C++. We set an initial cell configuration as a 20×20 cell tile, which was generated from randomly distributed centroids with a mean cell area of *A*_0_. We adopted a free boundary condition. The discretized time-step was set to Δ*t* = 0.1, and the simulation was run until a configuration of cells became stable (t = 5,000). To implement a T1 transition, junctions shorter than 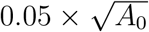 were allowed to perform a T1 transition if this transition reduces the virtual work.

To validate the parameter estimation using model A (section 3.1 and 3.5), we generated 15 synthetic data, whose parameters are set as follows. *λ*_0_ was fixed at 1.0, in line with the parameter estimation formulation, and *Γ* was fixed at 0.0. The other parameters were randomly generated from a logit-normal distribution. For instance, we set the distribution function of *λ*_1_ as follows:

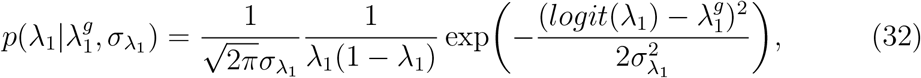

where 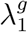 is the location parameter and 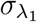 is the scale parameter for the group. The distribution function of the other parameters were set in the same way. The location parameters of the logit-normal distribution were set at 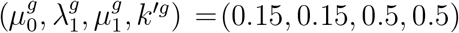, where *k*^*′*^ = 0.1*k*. The scale parameter was set at 0.3 for all parameters.

To validate the parameter estimation using model A’ (section 3.4), we generated 8 synthetic data using the following parameter sets: (*µ*_0_, *λ*_1_, *µ*_1_, *Γ, k*) = (0.15,0.12,0.5,0.04,4), (0.15,0.12,0.5,0.08,4), (0.05,0.15,0.5,0.05,5), (0.1,0.15,0.5,0.05,5), (0.1,0.15,0.5,0.1,5), (0.1,0.15,0.5,0.15,5), (0.2,0.15,0.5,0.05,5), (0.2,0.15,0.5,0.15,5) with *λ*_0_ being fixed at 1.0.

### 2.9 *In vivo* image data

*In vivo* image data used in this study have been described in [27]. *Drosophila* lines used were *sqhp-sqh-GFP* [54], *UAS-Dα-cat-TagRFP* [31], *apterous (ap)-Gal4*, and *DE-cad-GFP* [55]. The fly genotypes were *sqhp-sqh-GFP, ap-Gal4* /*sqhp-sqh-GFP, UAS-Dα-cat-TagRFP* for experiments in pupal wing and notum, and *DE-cad-GFP* for embryo. The total number of image data (samples) is 165. For the pupal wing, the number of image data at each developmental stage is as follows: 20 at 13-14 h APF, 23 at 16.5-18.5 h APF, 16 at 21-23 h APF, 21 at 25.5-27.5 h APF, and 25 at 30-32 h APF. For the pupal notum, there are 10 image data at each of five developmental stages (13-14 h, 16.5-18.5 h, 21-23 h, 25.5-27.5 h, and 30-32 h APF). For the embryo, there are 5 image data at each of two developmental stages (before and during the germband elongation).

### 2.10 Image processing and analysis

Images of the pupal wing and notum were segmented by custom-made macro and plugins in ImageJ/Fiji [31]. Images of the embryonic germband were segmented by EPySEG [56]. Vertex position and connectivity were extracted from skeletonized images by using custom-made code in OpenCV [31].

We preprocessed input image data as reporeted in [27]. Briefly, we excluded junctions shorter than 3 pixels (*in vivo* data) or 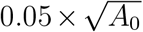 (synthetic data; *A*_0_ represents a target cell area) from the following analysis due to difficulties in measuring their angles. We also excluded cells larger than twice the median area of cells, since significantly larger cells, including sensory cells in the notum, could possess different mechanical properties from other cells. We then normalized the spatial scale using the median cell area. Finally, we rotated the vertex coordinates so that the body axes of each image taken from the same tissue were aligned (Fig. S2). We aligned the pupal wing images along the proximal-distal axis, while the pupal notum and embryonic germband images were aligned along the anterior-posterior axis. This step is necessary because pooled and hierarchical Bayesian estimation require the alignment of the body axis prior to conducting parameter estimation. Note that in our previous study [27], the axial parameters were corrected post-estimation rather than during the pre-processing of image data.

### 2.11 Bayesian force/stress inference

Here, we briefly describe the principle of Bayesian force/stress inference [31]. Bayesian force/stress inference solves an ill-conditioned, inverse problem between forces (junction tension and cellular pressure; denoted as 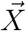in section 2.2) and cell shape. It derives the posterior distribution of cellular forces by combining the likelihood from the force-balance equation at cell vertices (Eq. 10) with the prior informed by experimental observations. The MAP estimate of 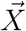 is obtained by maximizing the marginal likelihood function. Among various priors tested, the one assuming the positivity of junction tension has been shown to yield most accurate and robust estimates of the cellular forces [31, 41, 42, 57]. Details of the formalism, along with results of *in silico* and *in vivo* validation, have been reported in [31, 35, 41, 58–60].

## 3 Results

### 3.1 Validation of non-hierarchical Bayesian parameter estimation using synthetic data

The primary aim of this study is to extend and improve the image-based parameter inference [27] by reformulating it in a framework of Bayesian statistics. Image-based parameter inference involves the construction of models and the subsequent inference of parameters (Fig. 1D). For model construction, we followed the approach outlined in our previous study (section 2.1) [27]. The model equations for junction tension (referred to as the anisotropic spring model) and cell pressure were constructed based on image data of *Drosophila* epithelial tissues. The anisotropic spring model comprises two terms, Λ_0_(*θ*_*ij*_) and Λ_1_(*θ*_*ij*_)*l*_*ij*_ (Fig. 2 and Fig. S1). Λ_0_(*θ*_*ij*_) represents the line tension, whose magnitude is determined by the baseline *λ*_0_ with the anisotropic component (*µ*_0_, *φ*_0_). Similarly, Λ_1_(*θ*_*ij*_) consists of the baseline *λ*_1_ and an anisotropic component (*µ*_1_, *φ*_1_), which governs the dependency of junction tension on the junction length. In this and previous studies, *λ*_0_ was fixed at 1.0 as the scaling factor (section 2.2). We employed the standard model with area elasticity coefficient *k* for cell pressure (section 2.1). For parameter inference, by integrating non-informative prior distributions of these parameters with the likelihood function derived from the force balance at cell vertices, we established the non-hierarchical Bayesian parameter inference framework (section 2.4; Fig. 1G). The posterior distribution of parameters was computed through MCMC sampling (section 2.5).

To validate this newly developed non-hierarchical Bayesian method, we evaluated the accuracy of parameter estimation using synthetic data with known true parameter values. Synthetic data of the two-dimensional cell configuration was generated through numerical simulation of the CVM, as outlined in section 2.8 (Fig. 3A). Using the vertex position and connectivity in synthetic data as input, we obtained the posterior distribution and MAP estimates of the parameters. The estimation error was negligible, with the maximum error in relative value being 1.50% (Fig. 3B-E). These data clearly indicate that our non-hierarchical Bayesian method gives accurate estimates of the mechanical parameters *in silico*.

**Figure 3:**
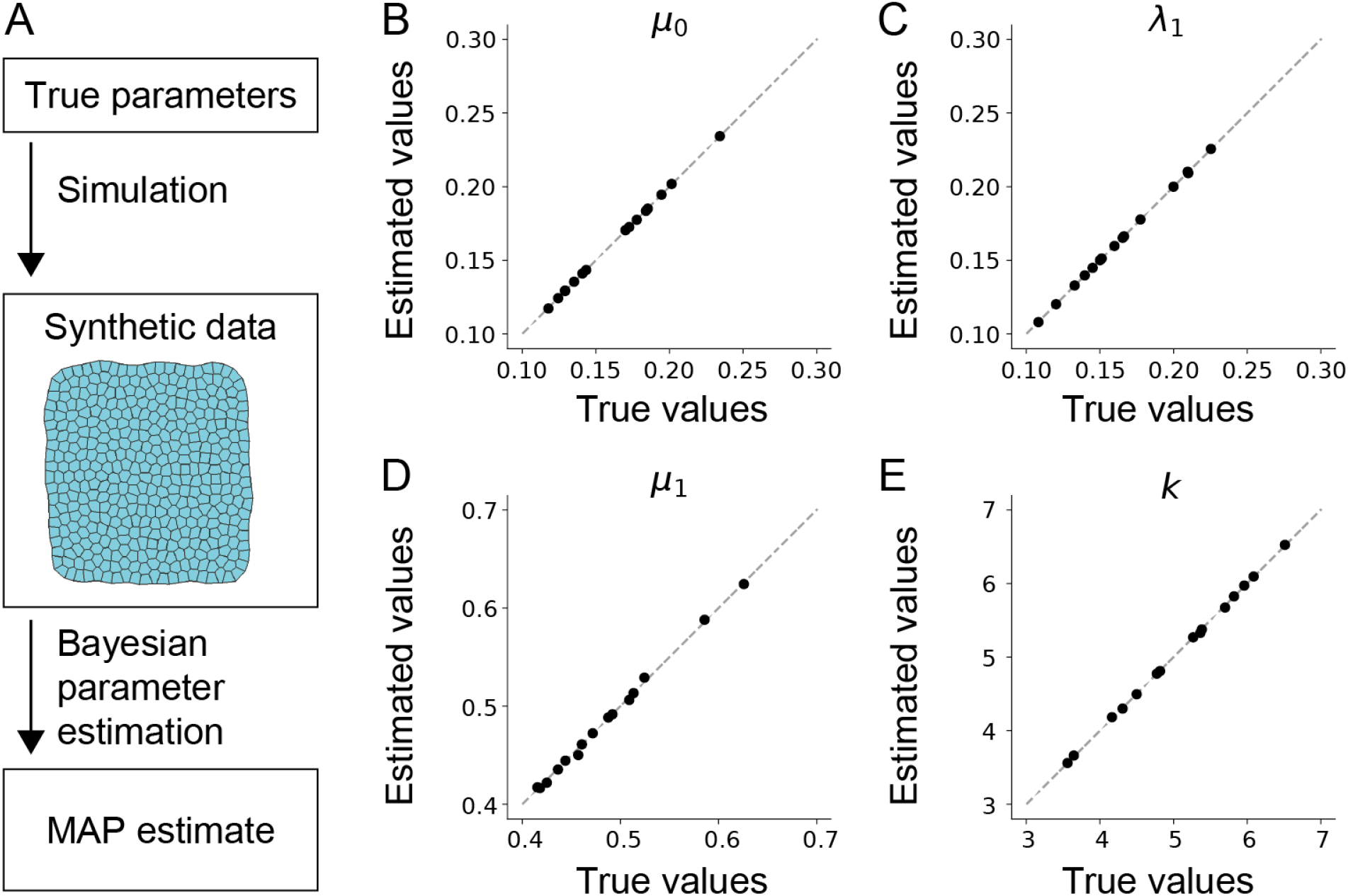
Validation of of the non-hierarchical Bayesian method using synthetic data. (A) Schematic of a perfect model experiment to asses the accuracy of parameter estimation. (B–E) Estimated values of parameters are plotted against their true values for *µ*_0_ (B; the magnitude of anisotropy in the line tension), *λ*_1_ (C; the spring constant of junction), *µ*_1_ (D; the magnitude of anisotropy in the spring constant of junction), and *k* (E; the area elasticity of cell). A dashed line indicates *y* = *x*. (A) is modified from [27].

### 3.2 Non-hierarchical Bayesian method yields parameter estimates consistent with those obtained by the least-squares method *in vivo*

Next, we evaluated the non-hierarchical Bayesian method on *in vivo* data. For this, we assessed its consistency with the least-squares method [27], which is generally expected when employing non-informative priors in a non-hierarchical Bayesian model [4]. Specifically, we applied the non-hierarchical Bayesian method to 165 image datasets of *Drosophila* epithelial tissues (section 2.9), conducted parameter estimation using the best-fitting model selected by the least-squares method, and then compared the estimated values between the two methods. Using the model selected by the least-squares method, the posterior distributions converged for all image data (samples), confirming the reliability of MCMC sampling. The MAP estimates were found to closely match the values inferred by the least-squares method, with a maximum difference in relative value of 1.65 × 10^*−*5^ for linear parameters, and a maximum difference in absolute value of 1.84 × 10^*−*6^ rad for axial parameters (Fig. 4A–F and Fig. S3).

**Figure 4:**
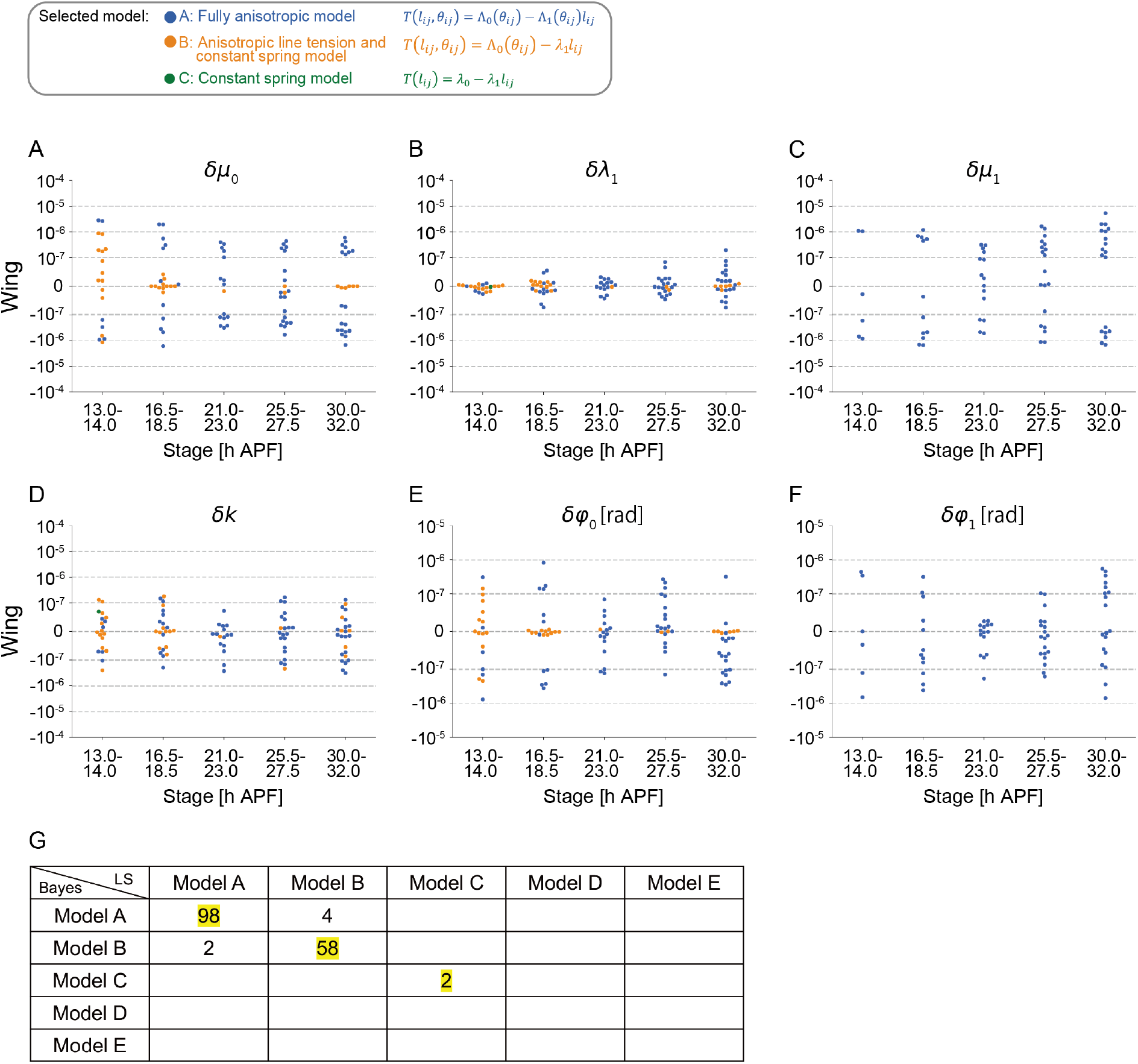
Comparison between the non-hierarchical Bayesian method and the original least-squares method using *in vivo* data. (A-F) The evaluation of parameter estimation of *µ*_0_ (A), *λ*_1_ (B), *µ*_1_ (C), *k* (D), *φ*_0_ (E), and *φ*_1_ (F). Each dot in (A-D) and in (E, F) respectively represents the relative difference and absolute difference in parameter estimations between the non-hierarchical Bayesian model (this study) and the original least-squares method [27] for each image data (sample) from the different developmental stages of the pupal wing. Dot colors represent the models selected by AIC in the original least-squares method, which were subsequently used for parameter estimation in the non-hierarchical Bayesian method. (G) Comparison of model selection between the original least-squares and non-hierarchical Bayesian methods. Each row and column represents the selected model by the original least-squares (LS) and the non-hierarchical-Bayesian (Bayes) methods, respectively. The numbers in the diagonal cells represent the number of images where the same model was selected by the LS and the Bayesian methods. MCMC sampling converged 823 out of the 825 total instances (165 image data across 5 candidate models).

We further assessed the consistency of model selection between the least-squares method and the non-hierarchical Bayesian method. The former selects models based on AIC [27], while the latter employs the widely applicable information criterion (WAIC) and leave-one-out cross-validation (LOOCV), which are commonly used model selection criteria in Bayesian statistics [49–51, 61] (section 2.6). We found that among candidate models A–E listed in Fig. S1, both WAIC and LOOCV consistently selected the same model for each image data. Using the model selected by WAIC and LOOCV, we could not obtain a converged posterior distribution for one image, which was subsequently excluded from further analysis. Our data indicated that the selected models were identical in the original and new methods for 158 out of the 164 images (Fig. 4G). For the remaining six images, the values of the model selection criteria were nearly indistinguishable between the rank-1 and rank-2 models, suggesting that the inconsistency in model selection arose from subtle variations in the model selection criteria (Appendix 2 in Supporting information). The fact that all three model criteria selected model A for most image data suggests that the anisotropy in positive feedback between junction tension and shrinkage plays important roles in *Drosophila* epithelial tissues. Overall, the tests on synthetic and *in vivo* data sets indicates that the image-based parameter inference has been adequately reformulated using Bayesian statistics.

### 3.3 Quantification of uncertainty in parameter estimation revealed its link to mechanical anisotropies in tissue

Assessing the uncertainty in parameter estimation offers insights into the reliability of obtained results [29]. In this study, we used 95% Highest Posterior Density Interval (HDI), the range within which a parameter value fall with a probability of 0.95, as a quantitative measure for the uncertainty associated with parameter estimation (Fig. 1F; section 2.7). Fig. 5A presents the 95% HDI values quantified for each parameter at each developmental stage in *Drosophila* pupal wing, in which different colors represent different models selected by WAIC. The 95% HDI values are shown as relative values for linear parameters and absolute values for axial parameters. The moderately larger 95% HDI of *µ*_1_ may be attributed to the higher order of its explanatory variable, *i*.*e*., the product of *l*_*ij*_ and cos 2(*θ*_*ij*_ *− φ*_1_). This could also explain why the 95% HDI values were smaller for simpler models.

**Figure 5:**
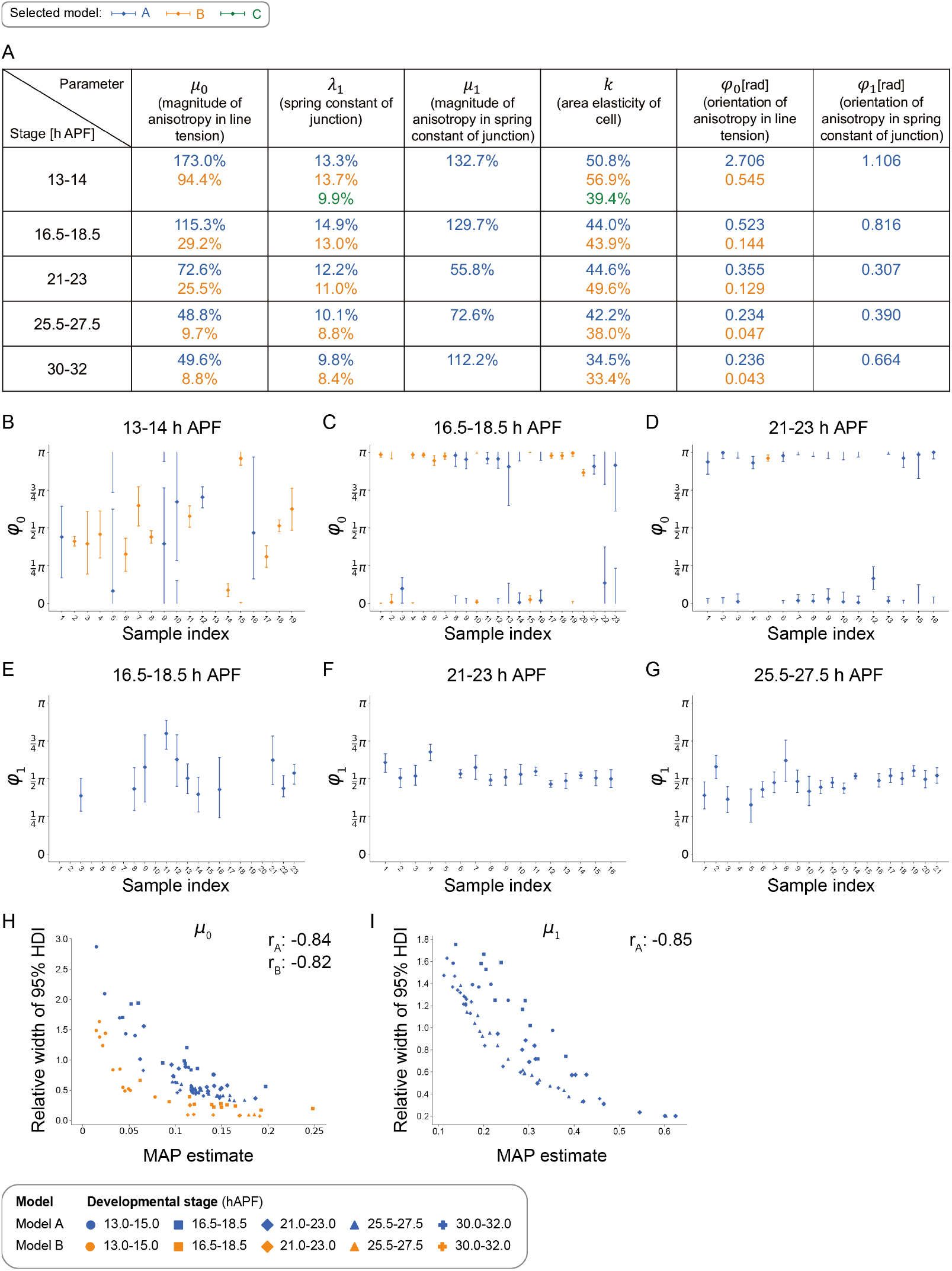
Quantification of uncertainty in parameter estimation using the non-hierarchical Bayesian method with *in vivo* data. (A) The average width of 95% HDI (section 2.7) is listed for each parameter at each developmental stage of the pupal wing. The 95% HDI values are normalized by the MAP estimate for linear parameters and are shown as absolute values for axial parameters. Each row corresponds to a developmental stage, indicated by hours after puparium formation (h APF), and each column corresponds to a model parameter. The colors represent the models selected by WAIC and LOOCV. (B-G) The 95% HDI for *φ*_0_ (B–D) and *φ*_1_ (E–G) at 13–14 (B), 16.5–18.5 (C, E), 21–23 (D, F) and 25.5–27.5 (G) h APF in the pupal wing. Bars and dots indicate the 95% HDI and the MAP estimate, respectively. Since we analyzed still images, the indices at different developmental stages are independent of each other. The indices for different parameters at each developmental stage correspond to the same image. (H, I) The correlation between the MAP estimate and the width of 95% HDI relative to the MAP estimate for *µ*_0_ (H) and *µ*_1_ (I) across all developmental stages of the pupal wing. The correlation coefficient for each selected model is shown at the top-right.

We then closely examined the 95% HDI values for individual parameters, focusing on their dependence on developmental stages. Our previous study has uncovered the distinctive functions of the tension parameters in the *Drosophila* pupal wing [27]: *µ*_0_ and *φ*_0_, representing the anisotropy in line tension, contribute to resistance against the proximal-distal (PD) tissue stretching induced by extrinsic forces (Fig. 2C, D). On the other hand, *µ*_1_ and *φ*_1_, representing the anisotropy in feedback between junction shrinkage and tension, facilitate the junction shrinkage along the anterior-posterior (AP) direction, perpendicular to tissue stretching (Fig. 2B, E). Here, we found that the 95% HDI of *φ*_0_ became narrower after 16.5 h APF, consistent with the timing of the onset of tissue stretching (Fig. 5A–D). The 95% HDI of *µ*_0_ exhibited a similar trend (Fig. 5A). These results suggest that as the wing begins to stretch, the estimation of *µ*_0_ and *φ*_0_ becomes more robust, reflecting the stabilization of tissue-scale mechanical structure. The 95% HDI of *µ*_1_ and *φ*_1_ took smaller values around the AP direction at 21–23 hr APF and 25.5–27.5 h APF, consistent with a stronger need to elevate the junction tension at remodeling AP junctions in these developmental stages (Fig. 5A, E-G). Other two parameters, *k* and *λ*_1_, did not exhibit significant changes in the 95% HDI values during the wing development (Fig. 5A, Fig. S4, Fig. S5). When analyzing data from all developmental stages together, the 95% HDI of *µ*_0_ and *µ*_1_ exhibited a negative correlation with their MAP values, corroborating the claim that estimation uncertainty decreases as the mechanical anisotropies increase (Fig. 5H, I). In summary, the results of uncertainty quantification revealed that the reliability of parameter estimation is tightly linked to the mechanical anisotropies present in the tissue.

### 3.4 The cortical elasticity term is dispensable for explaining force-shape correlations *in vivo*

Building on the non-hierarchical Bayesian method with non-informative priors, we further extended the image-based parameter inference by adding an informative prior on the cortical elasticity (section 3.4) and by constructing a hierarchical model (sections 3.5 and 3.6).

In the standard model function of the CVM, *T* (*L*_*i*_, *L*_*j*_) = *λ*_0_ +*Γ*(*L*_*i*_ +*L*_*j*_) (Eq. 5), the term *Γ*(*L*_*i*_ +*L*_*j*_) mathematically represents the elasticity of the cell cortex, where *Γ* is the elastic coefficient of the cell cortex, and *L*_*i*_ denotes the perimeter length of the *i*-th cell [15, 16, 22]. In this study, we statistically evaluated the impact of the cortical elasticity term on the mechanical properties of epithelial cells, which are characterized by force-shape correlations. We first analyzed the correlation between junction tension and cell perimeter length to see if they were positively correlated as the cortical elasticity term suggests (Fig. 6A). Despite considerable variation in cell perimeter length among wing cells, no significant correlation with junction tension was observed, suggesting a negligible influence of the cortical elasticity term on the variation in tension magnitude among junctions.

**Figure 6:**
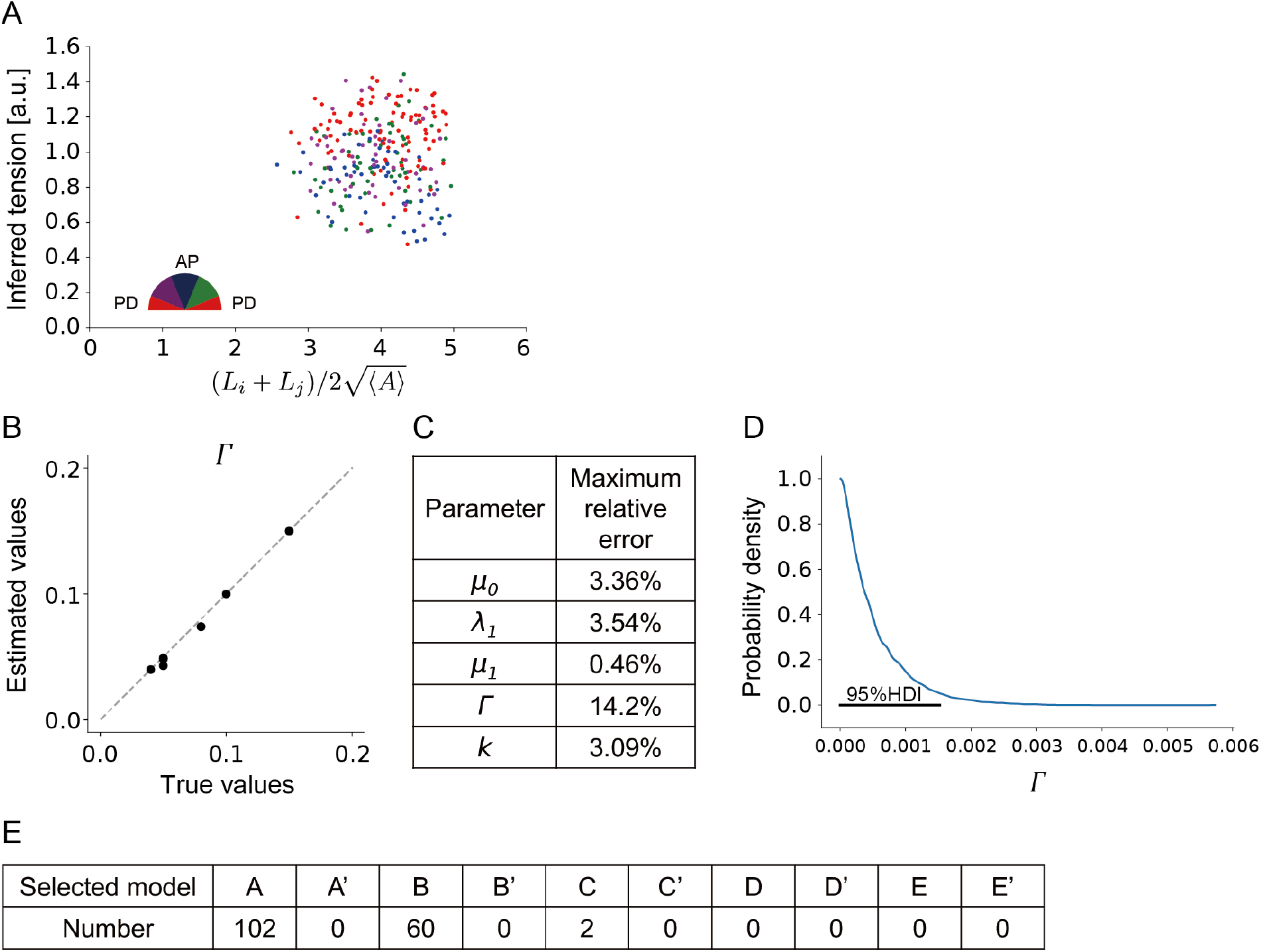
Parameter inference in the non-hierarchical Bayesian method that incorporates the cortical elasticity term. (A) The correlation between the inferred junction tension and the cell perimeter in *Drosophila* pupal wing at 18 h 5 min APF. Junction tensions are quantified using Bayesian force inference [31]. Each junction is classified by its orientation relative to the horizontal axis (semicircle; PD: proximal-distal and AP: anterior-posterior). The horizontal axis represents the average of the perimeters of cells adjoined by the same junction, normalized by the square root of the average cell area in the sample. (B, C) Accuracy of parameter estimation on synthetic data generated by numerical simulation of model A’ (Fig. S1B). (B) Estimated values of the parameter representing the elastic coefficient of cell cortex, *Γ*, in model A’ are plotted against their true values. (C) Table showing the maximum relative error in parameter estimation. (D) The posterior distribution of *Γ* obtained from the imaga data (sample) analyzed in (A). (E) Results of model selection between the anisotropic spring model with (A–E) and without (A’–E’) the cortical elasticity term (see Fig. S1 for the list of models). MCMC sampling converged 1644 out of the 1650 total instances (165 image data across 10 candidate models).

To support this finding, we sought to examine a model described by Eq. 4, which integrates the cortical elasticity term into the anisotropic spring model (model A’ in Fig. S1), and compared models with and without this term for its ability to fit to experimental data. However, with the original least-squares method, parameter estimation in model A’ frequently led to physically unrealistic, negative estimates for *Γ*. We therefore decided to adopt the non-hierarchical Bayesian method, imposing a prior that assumes the positivity of *Γ* (see section 2.4.1 for details). This extension enabled us to perform parameter estimation and model selection using model A’ and its simpler counterparts. When synthetic data was generated by the numerical simulation of model A’, both the WAIC and LOOCV consistently selected model A’ as the best data-fitted model among models A-E and A’-E’. Moreover, the estimated values closely aligned with the true values for all parameters, including *Γ* (Fig. 6B, C). These data clearly indicate that our method can properly handle models incorporating the cortical elasticity term. We then applied the extended non-hierarchical Bayesian method to the in vivo datasets. We found that the estimated values of *Γ* were close to zero (Fig. 6D), consistent with the minimal correlation between junction tension and cell perimeter length. Moreover, both the WAIC and LOOCV selected models in the category of the anisotropic spring model over those including the cortical elasticity term (Fig. 6E). In summary, all of our data suggest that the cortical elasticity term is not essential for explaining force-shape correlations *in vivo*.

### 3.5 Validation of hierarchical Bayesian parameter estimation using synthetic data

Classifying samples into groups, rather than treating them individually, and quantifying the characteristics of each group, provides valuable information for understanding the system of interest. Hierarchical Bayesian modeling offers a statistical framework that integrates individual variance with overall group properties, thereby yielding parameters at both the individual and group levels and identifying a most appropriate model of the group [62, 63]. To this end, we extended our Bayesian model into a hierarchical Bayesian model to estimate cell mechanical properties at both the group and individual levels (section 2.4.2; Fig. 1H). Specifically, we adopted the beta distribution as a prior for linear parameters to represent the variance among individual samples within the group. The half-Cauchy distribution is used for a prior for two parameters of the beta distribution, serving as the hyper prior. For axial parameters, we utilized the axial von Mises distribution, setting location parameters as the axial uniform distribution and concentration parameters as the reciprocal of a half-Cauchy distribution with a scale of 5.

We assessed the precision of parameter estimation of the hierarchical Bayesian method using synthetic data. Synthetic data was generated in the following procedures (Fig. 7A; section 2.8). We set true distributions to each parameter of model A, and generated random numbers from these distributions to create 15 sets of true parameters. Numerical simulation of the CVM was then carried out to obtain the polygonal cell configuration. Using the vertex positions and connectivity from the synthetic data as input, we conducted parameter estimation and obtained the posterior distributions at both the group (Fig. 7B) and the individual levels (Fig. 7C). At the individual level, the true and estimated parameter values closely matched (black dots in Fig. 7D–G). Estimation accuracy was high, evidenced by a maximum relative error of 1.52%. Furthermore, the MAP estimates of the group-level parameters were highly close to the mode of the true distribution (orange squares in Fig. 7D-G). For comparison, we aggregated the input data into a single dataset. The results from the pooled estimation deviated substantially from the true distributions (blue diamonds in Fig. 7D-G), presumably because the distinct characteristics of each input data were averaged out during the aggregation process. Collectively, our data highlight the effectiveness of hierarchical Bayesian modeling.

**Figure 7:**
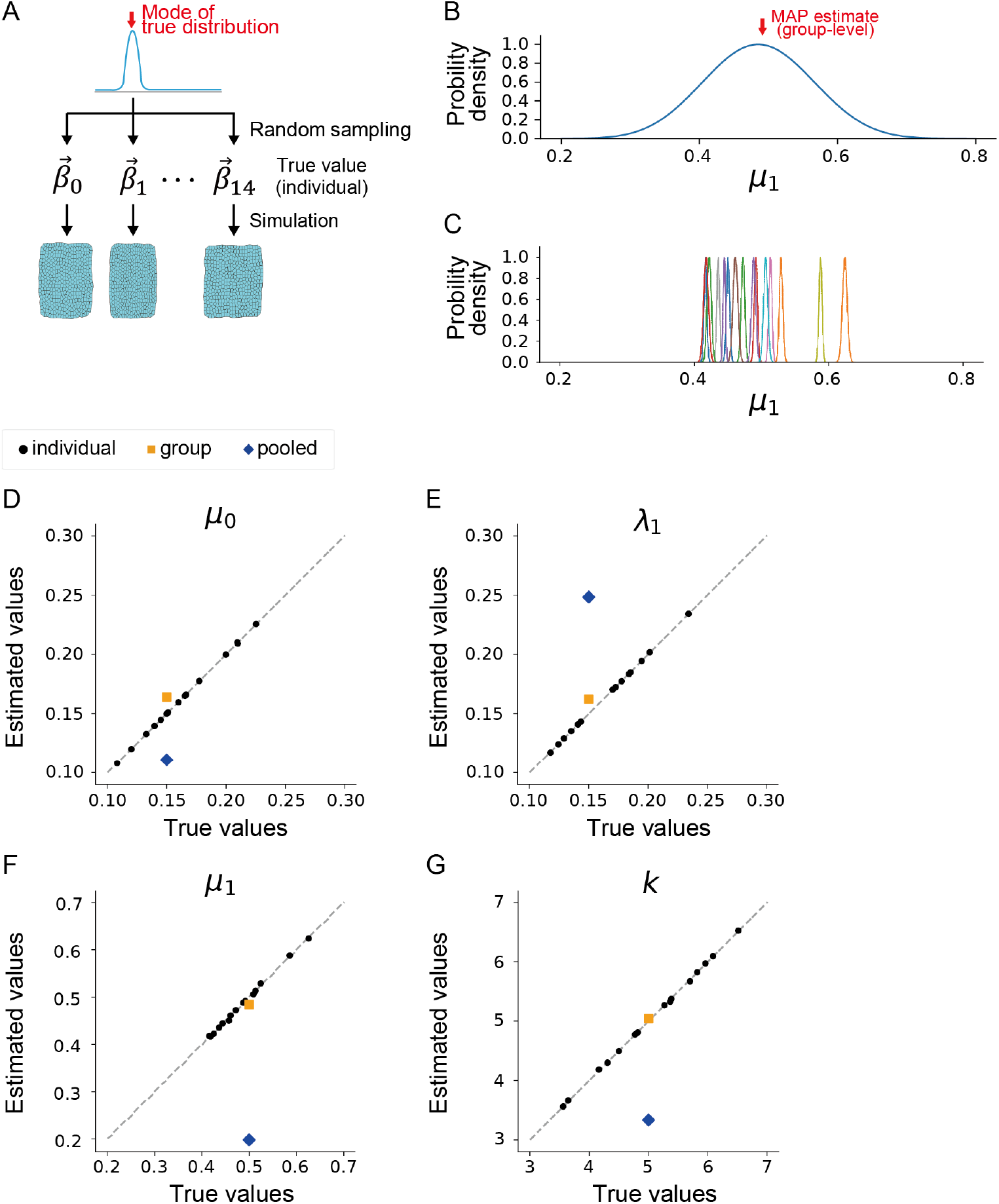
Validation of the hierarchical Bayesian model using synthetic data. (A) Schematic diagram illustrating the generation of synthetic data for testing the hierarchical Bayesian model. Individual-level parameters 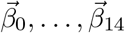 are sampled from the predefined group-level distribution. Synthetic data are then generated by numerically simulating the CVM using these parameters. (B, C) Posterior distribution of the group-level (B) and individual-level (C) parameters for *µ*_1_. In (C), different colors indicate different samples from the same developmental stage. (D–G) Estimated values of parameters are plotted against their true values for *µ*_0_ (D), *λ*_1_ (E), *µ*_1_ (F), and *k* (G). The black dots and orange squares represent the individual-level and group-level parameter estimates, respectively, obtained by the hierarchical Bayesian method. The blue diamonds show results from pooled estimation, where all individual data within a group are aggregated and then a single parameter set is estimated using the original least-squares method. A dashed line indicates *y* = *x*.

### 3.6 Application of hierarchical Bayesian method to *in vivo* data

Having established the effectiveness of our hierarchical Bayesian modeling, we applied the method to *in vivo* data from the *Drosophila* pupal wing. We conducted MCMC sampling across five developmental stages and for five candidate models A–E to obtain representative parameter values and models for each developmental stage. Our inference results indicated that the posterior distributions of mechanical parameters from all candidate models were effectively estimated at developmental stages starting from 16.5 h APF. The MCMC sampling did not converge for models A–C at 13–14 h APF, when cell and tissue mechanics do not exhibit strong anisotropies (see the Discussion for possible solutions). In the four developmental stages where MCMC sampling converged for all models, both WAIC and LOOCV selected the full model, model A, over its simpler counterparts. We then compared the parameter values estimated for model A by the hierarchical Bayesian method with those obtained by the least-squares method. We observed that compared to the results from the least-squares method, parameter values estimated by the hierarchical Bayesian method at the individual level showed a trend of shrinkage toward the group-level estimate (gray arrows in Fig. 8). This shrinkage could be attributed to the incorporation of a group-level distribution of parameters [29]. The group-level estimates aligned closely, regardless of the method used (green and orange dashed lines, and blue solid lines in Fig. 8), presumably because individual samples at the same developmental stage possessed similar parameter values.

**Figure 8:**
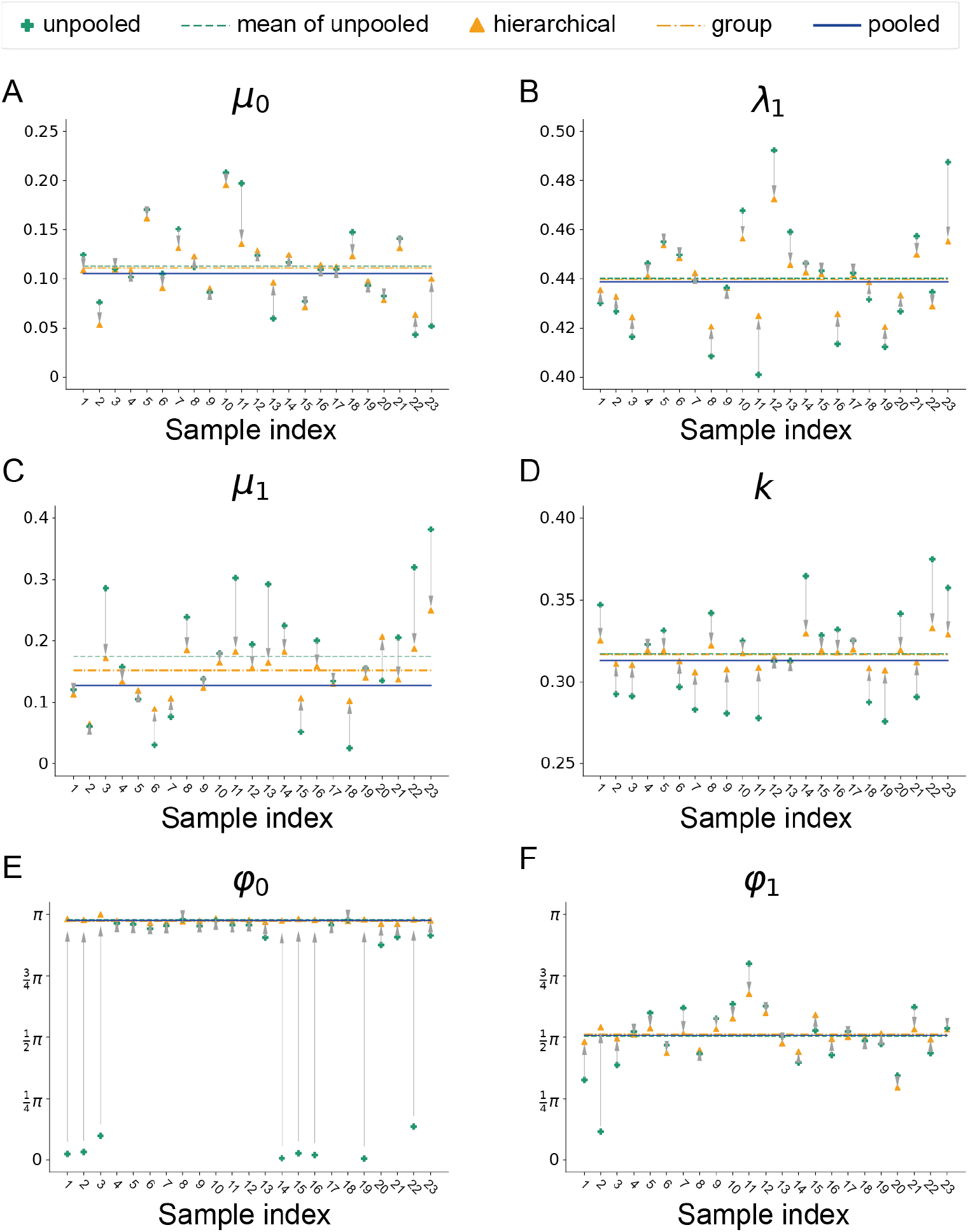
Application of the hierarchical Bayesian model to *in vivo* data. (A–E) Shrinkage plots for *µ*_0_ (A), *λ*_1_ (B), *µ*_1_ (C), *k* (D), *φ*_0_ (E), and *φ*_1_ (F) for data at the stage of 16.5–18.5 h APF in *Drosophila* pupal wing. Estimated values using the hierarchical Bayesian method are displayed at both the individual (orange triangles) and group levels (orange dashed line), alongside values estimated by the original least-squares method (green crosses), the average of these unpooled values (green dashed line), and the estimate from the pooled sample (blue solid line). Gray arrows highlight the shrinkage of estimated values from the original least-squares method to those obtained by the hierarchical Bayesian method.

Finally, we investigated how hierarchical Bayesian modeling influences estimation uncertainty. We analyzed the wing image data from 16.5–32 h APF, for which model A was selected by the non-hierarchical Bayesian method. Among these 64 images, the 95% HDI of all parameters took smaller values in the hierarchical Bayesian method for 51 images. For the remaining 13 images, the 95% HDI for some parameters increased; however, the difference was less than 10% in almost all cases. These results suggest that uncertainty in parameter estimation at the individual level decreases with hierarchical Bayesian modeling [61].

## 4 Discussion

To ensure the validity of mathematical modeling, selecting the most appropriate model among the candidates and determining its parameters based on experimental data is essential. By utilizing Bayesian statistics, this study extended the image-based parameter inference [27], a statistical framework designed for cell-based models in epithelial mechanics that computes model selection and parameter estimation. We demonstrated that Bayesian formulation enables the quantification of estimation uncertainty, broadens the range of models that can be analyzed, and facilitates the integration of information across different layers. These advantages significantly expand the utility of image-based parameter inference.

The uncertainty associated with parameter estimation has been assessed in only a limited number of studies involving cell-based mechanical models [7,8]. Our quantification of estimation uncertainty revealed significant changes in the 95% HDI values during wing development: the 95% HDI of anisotropy parameters in tension functions decreased concomitantly with the increase in the magnitude of anisotropy in cell/tissue mechanics. These findings strongly suggest that the estimation of mechanical parameters becomes more robust when cell/tissue mechanics are aligned across the tissue and the tissue-scale mechanical structure is stabilized. Such information helps researchers gauge the reliability of theoretical predictions.

Other factors can potentially affect the uncertainty in parameter estimation with our original least-squares and new Bayesian methods. Variance in force balance at the cell vertex, including violations of the quasi-static assumption, can directly impact the equations solved to infer parameters. Bayesian force/stress inference is employed solely for formulating candidate model equations and does not influence the parameter estimation itself. Measurement errors in vertex positions are likely of moderate relevance, as parameter estimation has been shown to be relatively robust to noise in vertex positions in our *in vivo* images [27]. However, this can depend on the quality of the image and image processing.

Our previous study revealed that the anisotropic spring model is favored over the standard CVM for fitting experimental data of *Drosophila* epithelial tissues [27]. However, the original least-squares method could not accommodate a model that integrates the cortical elasticity term with the anisotropic spring model. Bayesian modeling addressed this limitation by incorporating prior knowledge about the elasticity of the cell cortex. This reformulation enabled a comparison between the anisotropic spring model with and without the cortical elasticity term based on statistical criteria, as validated by the test using synthetic data. The model selection analysis revealed that for all *in vivo* data tested in this study, models excluding the cortical elasticity term were selected. These results suggest that the cortical elasticity term may not be essential for explaining the force-shape correlation that represent the cell mechanical properties. Alternatively, because there needs to be significant variation in the geometry among cells in the input image data for image-based parameter inference to work properly, the strong physical constraint on the cell perimeter length can hinder the detection of the cortical elasticity term. However, our measurement of cell geometry indicated significant variance in the perimeter length among cells, which contradicts the existence of such a strong physical constraint. Recent studies also highlight the need to revisit the standard equations in cell-based models in epithelial mechanics [17–21]. Some of these revised CVM equations do not include the cortical elasticity term; instead, they incorporate a time derivative of line tension that negatively depends on junction strain [18,20,21]. It is possible that extending image-based parameter inference to consider temporal dynamics could lead to the selection of models incorporating such time-dependent feedback.

In constructing the hierarchical Bayesian inference, we employed prior distributions based on the assumption that each parameter for a given developmental stage is centered around a specific value. While this hierarchical Bayesian model successfully estimated model parameters for most developmental stages, MCMC sampling became unstable at certain developmental stages where the parameters controlling the strength of tension anisotropy, specifically *µ*_0_ or *µ*_1_, were small. During these stages, the parameters governing anisotropy orientation, namely *φ*_0_ or *φ*_1_, showed significant variation among individual samples that the aforementioned prior distribution could not adequately capture, leading to the instability of MCMC sampling. Therefore, the convergence of MCMC sampling in the hierarchical Bayesian model exhibited a similar trend to estimation uncertainty in the non-hierarchical Bayesian model. We observed that MCMC sampling was stabilized upon adopting a uniform distribution as the prior for the concentration parameter of the axial Von-Mises distribution (*κ*), which controls the variation of *φ*_0_ and *φ*_1_. These results suggest the possibility of modifying the hyper prior distribution to incorporate the characteristics of the system of interest.

Adapting Bayesian parameter inference to handle various types of models and data will create opportunities for future research. For instance, by segmenting a whole-tissue image into smaller regions and applying tissue-level hyper-priors in hierarchical Bayesian modeling, one can infer a spatially smooth distribution of parameters while accounting for spatial variations within the tissue. Similarly, using time-lapse data as input, hierarchical Bayesian modeling can integrate both global trend in time-dependent changes and the variance among time-series of mechanical parameters. Furthermore, given recent advancements in cell-based modeling, especially those regarding feedback regulations in epithelial junction [18–20, 64, 65], analyzing recently proposed model functions by using the Bayesian parameter inference will be of great interest. These extensions will contribute to deepening our understanding of cell mechanics during tissue development and repair.

## Data availability

The authors declare that the data supporting the findings of this study are available within the paper and its Supplementary files. The code used for Bayesian parameter inference and input data used in this study can be downloaded from https://github.com/Sugimuralab/BayesianParameterInferenceForEpithelialMechanics and https://ssbd.riken.jp/repository/270/, respectively.

## CRediT authorship contribution statement

**Xin Yan**: formal analysis, investigation, methodology, software, writing - original draft; **Goshi Ogita**: conceptualization, project administration, formal analysis, investigation, methodology, writing - original draft, funding acquisition; **Shuji Ishihara**: conceptualization, supervision; writing - review and editing, funding acquisition; **Kaoru Sugimura**: conceptualization, project administration, supervision, writing - original draft, writing - review and editing, funding acquisition. All authors gave their final approval for publication and agreed to be held accountable for the work performed therein.

## Acknowledgments

The authors would like to thank Yang Hong, Roger Kares, the Bloomington Stock Center, and the Kyoto Stock Center for fly strains; and Mayu Miyakawa for technical assistance. X.Y. and G.O. are respectively supported by World-leading Innovative Graduate Study for Frontiers of Mathematical Sciences and Physics (WINGS-FMSP) and RIKEN Pioneering Project “Prediction for Science”.

## Supplementary information

## Appendix 1 Implementation of the likelihood function

In section 2.4.1, we defined the likelihood function as the probability density function of the n-dimensional multivariate normal distribution (Eq. 26). Since we assume the variance-covariance matrix to be *σ*^2^*I*, the likelihood function can be transformed into the product of the 1 -dimensional normal distribution as follows:

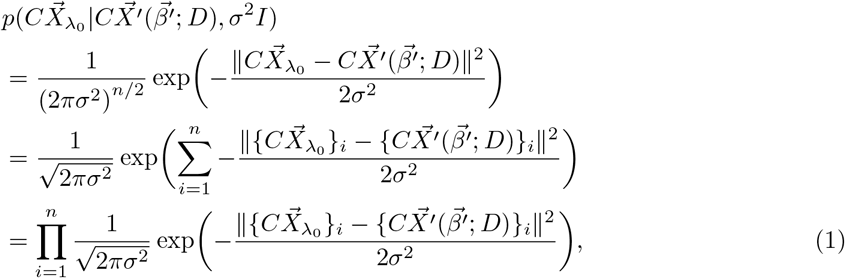

where 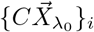 and 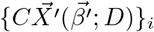 respectively represent the *i*-th component of 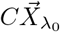 and 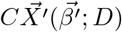 In the code used for Bayesian parameter inference (https://github.com/Sugimuralab/BayesianParameterInferenceForEpithelialMechanics), we implemented the likelihood function using the expression in the last row of the above equation.

## Appendix 2 WAIC values for rank-1 and rank-2 models

In Section 3.2, we compared the best-fit model selected by AIC in the original least-squares method with the model selected by WAIC in the new Bayesian methods, showing that the same model was chosen for 158 out of 164 image data of Drosophila epithelial tissues. For the remaining six images, the WAIC values of rank-1 and rank-2 models were as follows: (model A: 182.95, model B: 182.73), (model A: 592.68, model B: 592.64), (model A: 644.25, model B: 644.16), (model B: 571.36, model A: 571.24), (model A: 255.16, model B: 254.82) and (model B: 208.76, model A: 208.32), respectively. These results suggest that the inconsistencies in model selection arose from subtle variations in the model selection criteria.

**Figure S1:**
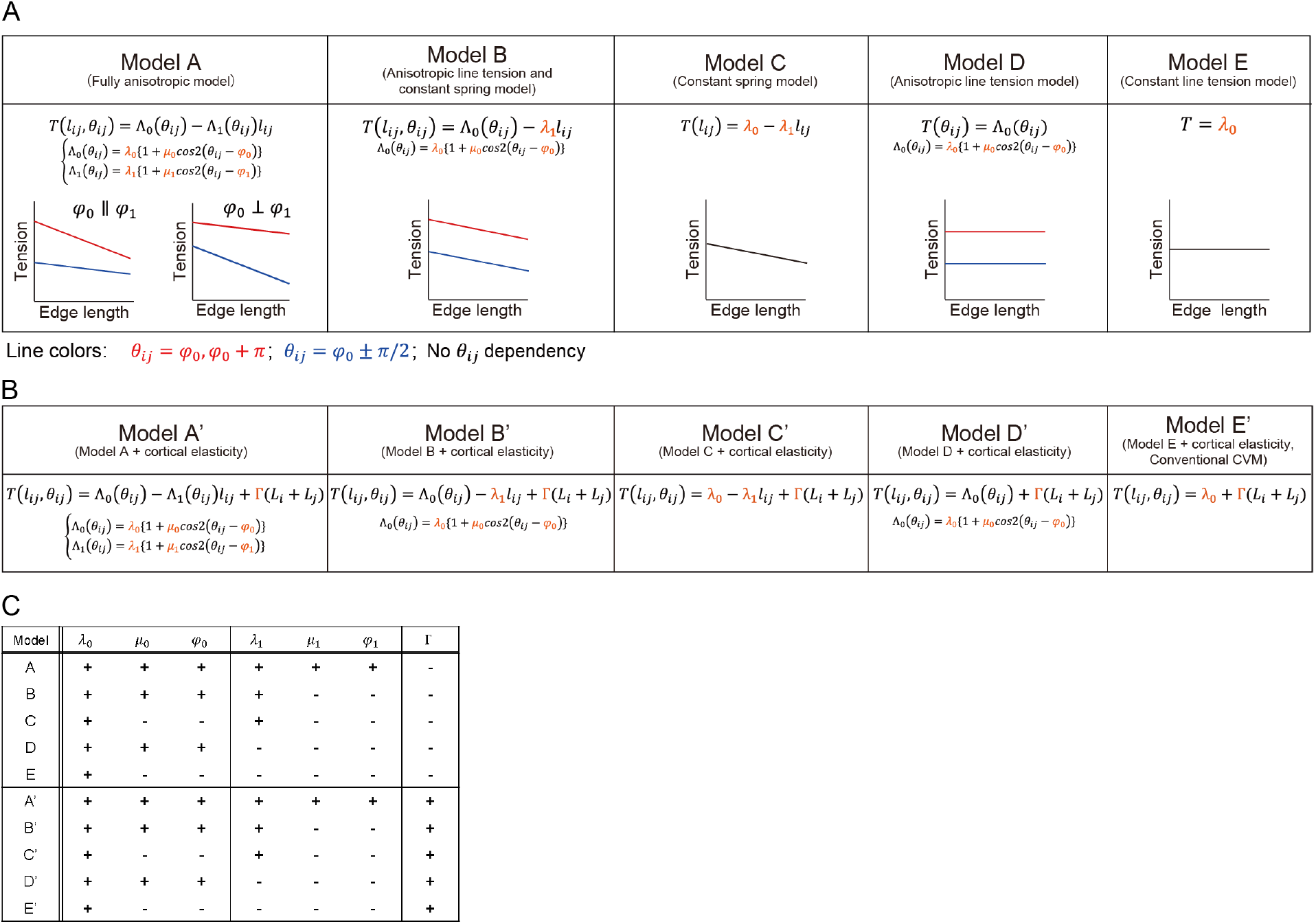
Candidate tension models. (A) Anisotropic spring model (model A) and its simpler counterparts (model B to E). Equations for junction tension are provided for each candidate model, with parameters highlighted in orange. Schematics show how the values of *µ*_0_ and *µ*_1_ influence the dependence of tension on junction length and orientation. Line colors indicate the orientation of the junctions. (B) Anisotropic spring model with cortical elasticity (model A’) and its simpler counterparts (model B’ to E’). Equations for junction tension are provided for each candidate model, with parameters highlighted in orange. (C) Table listing all candidate tension models used in this study. Each row corresponds to a candidate model, and each column to a parameter. The + and *−* signs respectively indicate whether a parameter was included or excluded from model equations. (A) is adapted from [27].

**Figure S2:**
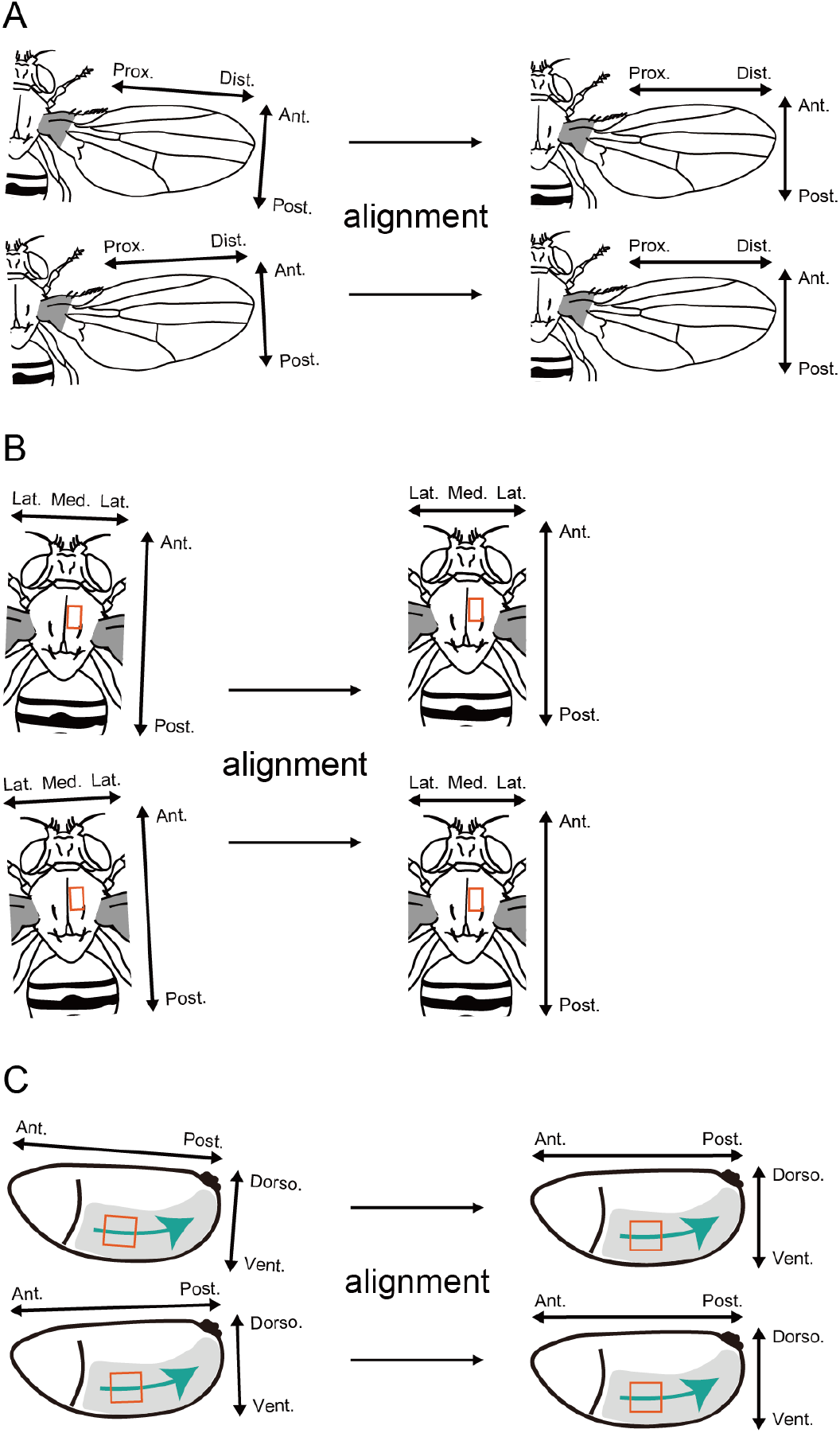
Alignment of *in vivo* data in image processing. (A–C) Schematics illustrating the alignment process for image data from the wing (A), notum (B), and embryonic germband (C). The original images on the left are rotated to align along the tissue axis. Adapted from [27].

**Figure S3:**
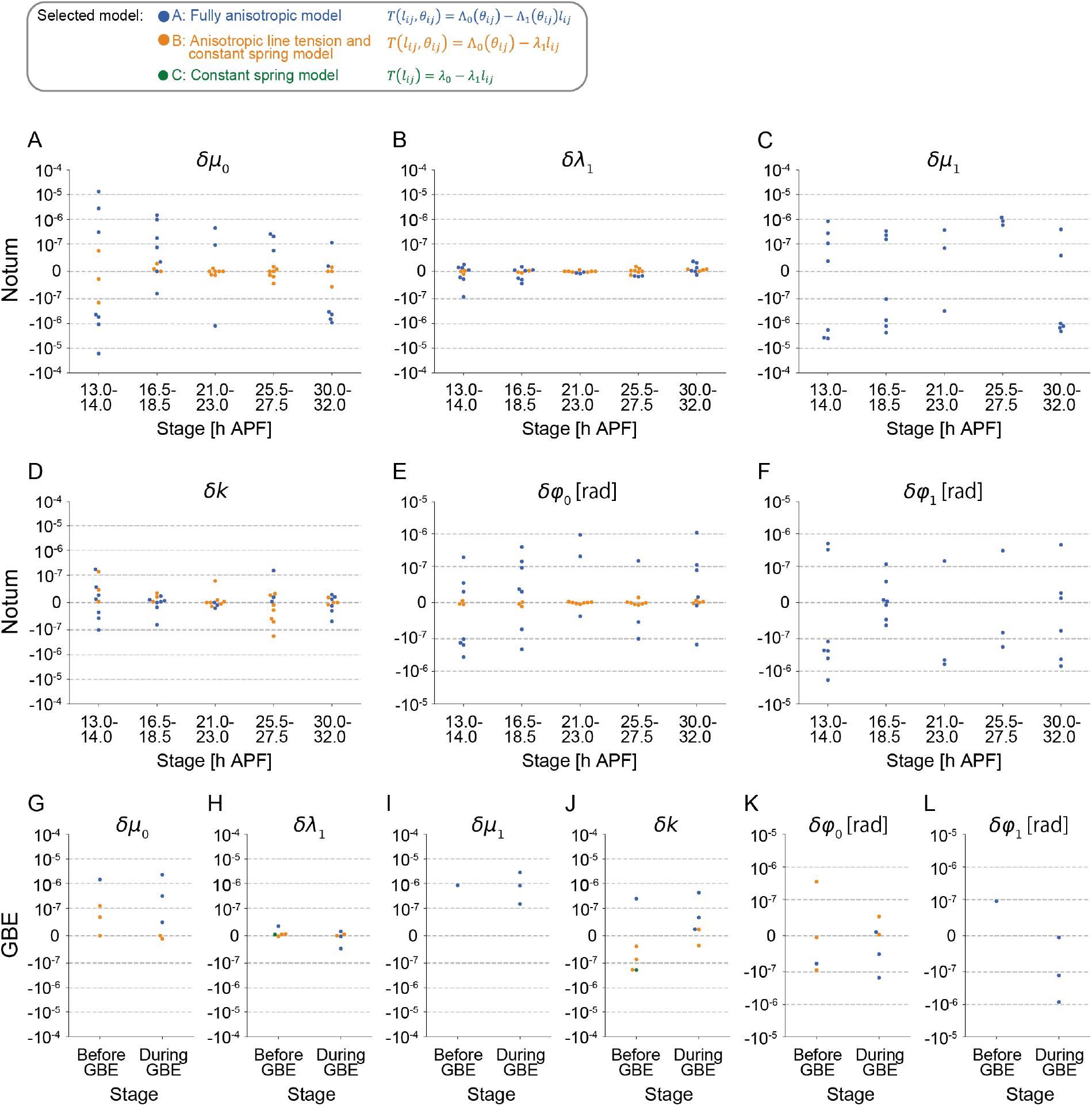
Comparison between the non-hierarchical Bayesian method and the original least-squares method using *in vivo* data. (A-L) Evaluation of parameter estimation for *µ*_0_ (A, G), *λ*_1_ (B, H), *µ*_1_ (C, I), *k* (D, J), *φ*_0_ (E, K), and *φ*_1_ (F, L). Each dot in (A-D, G-J; linear parameters) and (E, F, K, L; axial parameters) respectively represents the relative difference and absolute difference in parameter estimates between the non-hierarchical Bayesian method and the original least-squares method for each image (sample) from different developmental stages of the pupal notum (A-F) and embryonic germband (GBE) (G-L). Dot colors represent the models selected by AIC in the original least-squares method, which were subsequently used for parameter estimation in the non-hierarchical Bayesian method.

**Figure S4:**
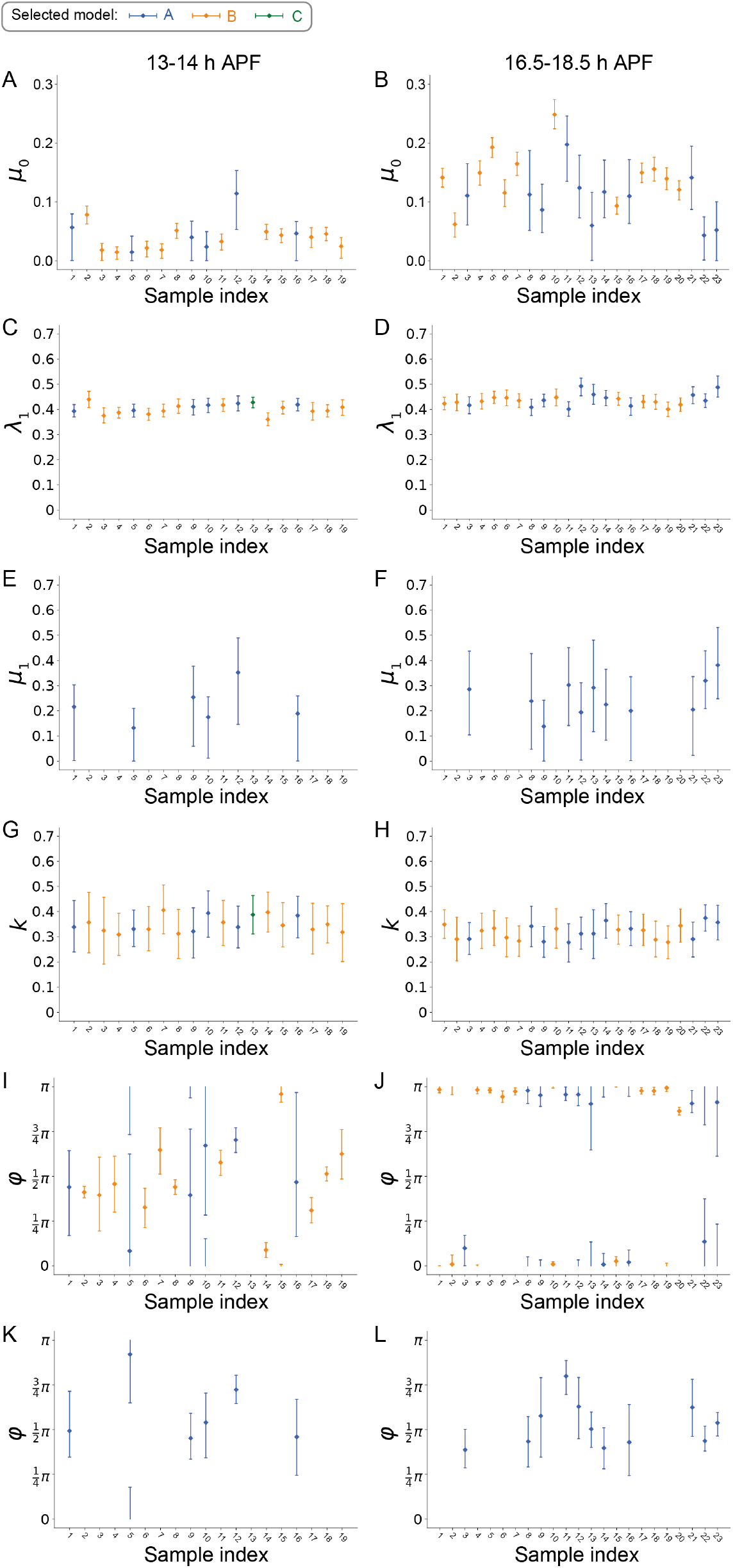
95% HDI of parameters estimated using the non-hierarchical Bayesian model with *in vivo* data. (A-L) The 95% HDI for parameters *µ*_0_ (A, B), *λ*_1_ (C, D), *µ*_1_ (E, F), k (G, H), *φ*_0_ (I, J), and *φ*_1_ (K, L) at 13-14 (A, C, E, G, I, K) and 16.5-18.5 (B, D, F, H, J, L) h APF in pupal wing. Bars and points indicate the 95% HDI and the MAP estimate, respectively. The index represents different image data from each developmental stage. The colors represent the models selected by WAIC and LOOCV.

**Figure S5:**
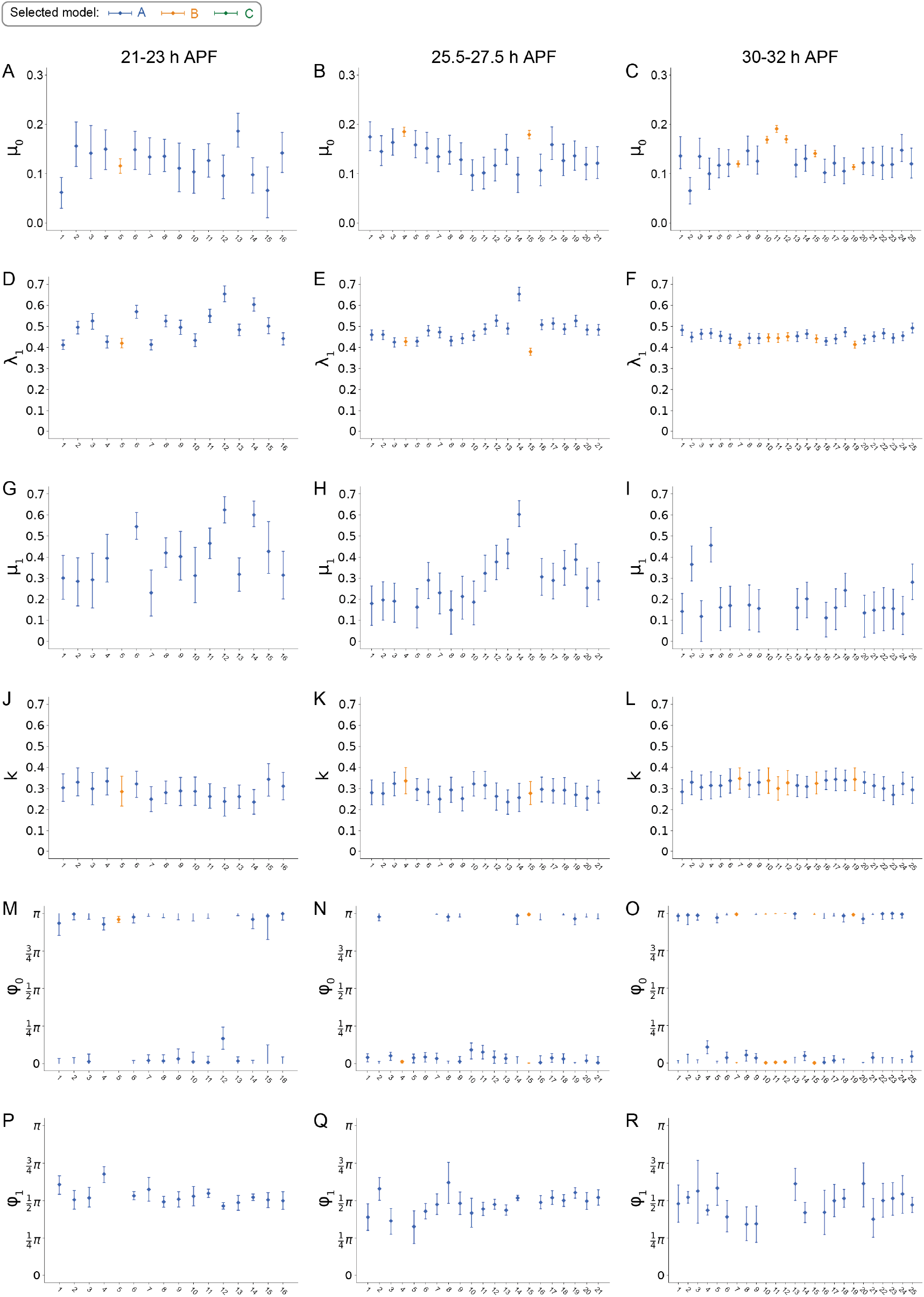
95% HDI of parameters estimated using the non-hierarchical Bayesian model with *in vivo* data. (A-R) The 95% HDI of the posterior distributions for parameters *µ*_0_ (A-C), *λ*_1_ (D-F), *µ*_1_ (G-I), k (J-L), *φ*_0_ (M-O), and *φ*_1_ (P-R) at 21-23 (A, D, G, J, M, P), 25.5-27.5 (B, E, H, K, N, Q) and 30-32 (C, F, I, L, O, R) h APF in pupal wing. Bars and points indicate the 95% HDI and the MAP estimate, respectively. The index represents different image data from each developmental stage. The colors represent the models selected by WAIC and LOOCV.

